# Neural Basis of the Sound-Symbolic Crossmodal Correspondence Between Auditory Pseudowords and Visual Shapes

**DOI:** 10.1101/478347

**Authors:** Kelly McCormick, Simon Lacey, Randall Stilla, Lynne C. Nygaard, K. Sathian

**Affiliations:** Department of Psychology, Emory University, Atlanta, GA 30322, USA; Winship Cancer Institute, Emory University, Atlanta, GA 30322, USA; Department of Neurology, Penn State College of Medicine, Hershey, PA 17033-0859, USA; Neural and Behavioral Sciences Penn State College of Medicine, Hershey, PA 17033-0859, USA; Psychology, Milton S. Hershey Medical Center, Penn State College of Medicine, Hershey, PA 17033-0859, USA

**Keywords:** multisensory, magnitude, semantic, phonology, attention, congruency effect

## Abstract

Sound symbolism refers to the association between the sounds of words and their meanings, often studied using the crossmodal correspondence between auditory pseudowords, e.g. ‘takete’ or ‘maluma’, and pointed or rounded visual shapes, respectively. In a functional magnetic resonance imaging study, participants were presented with pseudoword-shape pairs that were sound-symbolically congruent or incongruent. We found no significant congruency effects in the blood oxygenation level-dependent (BOLD) signal when participants were attending to visual shapes. During attention to auditory pseudowords, however, we observed greater BOLD activity for incongruent compared to congruent audiovisual pairs bilaterally in the intraparietal sulcus and supramarginal gyrus, and in the left middle frontal gyrus. We compared this activity to independent functional contrasts designed to test competing explanations of sound symbolism, but found no evidence for mediation via language, and only limited evidence for accounts based on multisensory integration and a general magnitude system. Instead, we suggest that the observed incongruency effects are likely to reflect phonological processing and/or multisensory attention. These findings advance our understanding of sound-to-meaning mapping in the brain.

## INTRODUCTION

Although the relationship between the sound and meaning of a word is often considered primarily arbitrary (de Saussure, 1916/2009; Pinker, 1999; Jackendoff, 2002; but see also Joseph, 2015), there is evidence that sound-meaning mappings occur reliably in a variety of natural languages (Blasi et al., 2016): such mappings are termed ‘sound-symbolic’ (e.g., Svantesson, 2017). A clear example of sound-meaning mapping is onomatopoeia, in which the sound of a word mimics the sound that the word represents (Catricalà and Guidi, 2015; Schmidtke et al., 2014), for example: ‘fizzle’, ‘bang’, or ‘splash’. The sound-symbolic relation in onomatopoeia is within-modal, i.e., word sounds are used to index sound-related meanings, but in other classes of words, sounds can index visual, tactile, or other sensory meanings, i.e., the sound-symbolic relation is crossmodal. For example, Japanese has a distinct class of words called mimetics including ‘*kirakira’* (flickering light) and ‘*nurunuru’* (the tactile sensation of sliminess: Akita and Tsujimura, 2016; Kita, 1997). Sound symbolism is often studied by examining the crossmodal correspondence between auditory pseudowords and two-dimensional visual shapes: e.g. ‘takete’ or ‘kiki’ are matched with pointed shapes whereas ‘maluma’ or ‘bouba’ are matched with rounded shapes (Köhler, 1929, 1947; Ramachandran and Hubbard, 2001). In this kind of audiovisual pairing, the shapes can be considered as representing potential referents or meanings for the pseudowords^1^. However, the conditions under which sound-symbolic associations arise are not clear. Spence (2011) proposed that crossmodal correspondences may be rooted in statistical, structural or semantic origins; these closely approximate explanations for sound-symbolic associations proposed by Sidhu and Pexman (2018).

One possible explanation for sound symbolism is that the relationships between sound-symbolic words and their visual or semantic referents reflect audiovisual statistical regularities in the natural environment that are integrated into a single perceptual object, for example, the sight of a stone thrown into water and the sound of it splashing might give rise to the onomatopoeic sound-to-meaning correspondence mentioned above. Such regularities may also underlie other crossmodal correspondences (Spence, 2011), for example that between high/low auditory pitch and high/low auditory and visual elevation (Jamal et al., 2017), since high-pitched sounds tend to emanate from high locations and low-pitched sounds from low locations (Parise et al., 2014).

Thus, sound symbolism might be related to a more general multisensory integration process in which auditory and visual features are linked (Ković et al., 2010) and which may also underlie other crossmodal correspondences (Sidhu and Pexman, 2018). If so, neural activity related to sound-symbolic processing might co-localize with activity related to multisensory integration, e.g. in the superior temporal sulcus (STS) when audiovisual synchrony (Beauchamp, 2005a,b; van Atteveldt et al., 2007; Stevenson et al., 2010; Marchant et al., 2012; Noesselt et al., 2012; Erickson et al., 2014) or audiovisual identity (Sestieri et al., 2006; Erickson et al., 2014) are manipulated, or in regions such as the intraparietal sulcus (IPS) when audiovisual spatial congruency is manipulated (Sestieri et al., 2006).

Sensory features can often be characterized along polar dimensions of magnitude where one end is ‘more than’ the other (Smith and Sera, 1992). Both visuospatial attributes of shapes (e.g., size, spatial frequency) and acoustic-phonetic attributes of speech sounds (e.g., sonority, formant frequencies) could be encoded by a domain-general magnitude system (Dehaene et al., 2003; Walsh, 2003; Lourenco and Longo, 2011) in which different attributes might become associated by virtue of occupying similar positions along magnitude dimensions. Such crossmodal magnitude relations have been proposed to underpin some crossmodal correspondences (Lourenco and Longo, 2011; Spence, 2011) as well as sound-symbolic associations (Sidhu and Pexman, 2018). Spence (2011) referred to this possibility as a “structural” account emerging as a consequence of modality-independent neural coding mechanisms, e.g. the neural coding of stimulus magnitude in neuronal firing rates across multiple sensory modalities. Thus, in sound-symbolic language, magnitude representations may serve to link the sound structure of spoken language with object shapes (which can vary along dimensions of pointedness, roundedness, or other shape attributes). That judgments of the sounds of language vary systematically along such dimensions is supported by findings that pseudowords with varying collections of phonetic features are readily placed along continua of pointedness and roundedness (McCormick et al., 2015), and that ratings transition from rounded to pointed with increasing vocal roughness, as indexed by a number of acoustic parameters (Lacey et al., 2020). On this account, activity related to sound symbolism might be expected in the intraparietal sulcus (IPS), a region involved in processing both numerical and non-numerical (e.g., luminance) magnitude (Sathian et al., 1999; Eger et al., 2003; Walsh, 2003; Pinel et al., 2004; Piazza et al., 2004, 2007; Sokolowski et al., 2017).

The third possible explanation for crossmodal correspondences proposed by Spence (2011) is semantic mediation. By this account, an unfamiliar word or pseudoword could be associated with a particular meaning by virtue of its similarity to other, real, words since crossmodal sound-symbolic associations exist in natural language, for example, ‘balloon’ and ‘spike’ for rounded and pointed shapes (Sučević et al., 2015; Blasi et al., 2016; Sidhu et al., 2021). Familiar words such as these have established semantic, as well as sound-symbolic, associations. Thus, when assigning ‘maluma’ to a rounded shape (Köhler, 1929, 1947), people might invoke real words with similar phonological content (for example, ‘balloon’). This semantic mediation account predicts that pseudoword-shape associations might activate regions of the left hemisphere language network.

Despite extensive behavioral investigations of sound symbolism, there have been relatively few studies of its neural basis and each has some limitations. Some studies employed a variety of sound-symbolic words and referent types (Ković et al., 2010; Revill et al., 2014; Sučević et al., 2015; Lockwood et al., 2016), which may obscure processing differences between different sound-symbolic effects (Sidhu and Pexman, 2018). Some earlier functional magnetic resonance imaging (fMRI) studies did not attempt to distinguish between potential explanations (Revill et al., 2014; Peiffer-Smadja and Cohen, 2019); whereas EEG studies (Ković et al., 2010; Sučević et al., 2015; Lockwood et al., 2016), while providing excellent temporal information, may not have sufficient spatial resolution to identify the anatomical sources of activity in the brain.

In the present study, we used fMRI to investigate cerebral cortical localization of differential activations for congruent vs. incongruent sound-symbolic crossmodal correspondences between auditory pseudowords and visual shapes. We chose to rely on implicit correspondences by presenting audiovisual pseudoword-shape pairs, asking participants to discriminate either the auditory pseudowords or the visual shapes. In order to investigate the competing explanations outlined above, we conducted three independent task contrasts in the same individuals, reflecting multisensory integration, magnitude estimation, and language processing. We hypothesized that a potential role for one of these accounts would be supported by finding cortical activations common to one of these independent task contrasts and the sound-symbolic (in)congruency between pseudowords and shapes. On the other hand, cortical activations lacking overlap with one of the independent task contrasts would require an alternative explanation. Finally, few studies have examined individual differences in crossmodal correspondences. We addressed this by administering the Object-Spatial Imagery and Verbal Questionnaire (OSIVQ: Blazhenkova and Kozhevnikov, 2009). We hypothesized that individuals with a preference for verbal processing might be more adept at assigning potential meanings to pseudowords; this hypothesis would be supported by significant correlations between task performance and/or neural activity with the verbal sub-scale of the OSIVQ. Additionally, object imagers might be more apt to visualize the shapes associated with pseudowords since such individuals typically integrate multiple sources of information about an object, compared to the more schematic representations of spatial imagers (Blazhenkova and Kozhevnikov, 2009; Lacey et al., 2011). This hypothesis would be supported by significant correlations between task performance and/or neural activity with preferences for object, rather than spatial, imagery.

## MATERIALS AND METHODS

### Participants

Twenty participants took part in this study, but one was excluded for excessive movement in the scanner (> 1.5mm), leaving a final sample of 19 (9 male, 10 female; mean age 25 years, 1 month). All participants were right-handed based on a validated subset of the Edinburgh handedness inventory (Raczkowski et al., 1974) and reported normal hearing and normal, or corrected-to-normal, vision. All participants gave informed consent and were compensated for their time. All procedures were approved by the Emory University Institutional Review Board.

### Procedures

#### General

Five participants took part in the pseudoword-shape scans first, and then underwent three scans to test competing accounts of the processes underlying the pseudoword-shape correspondence. The remaining participants had already undergone these latter scans approximately four months earlier as part of a separate study and thus completed the pseudoword-shape scans after these scans. After completing the required scan sessions, all participants performed a behavioral task to determine the strength of their crossmodal pseudoword-shape correspondence. All experiments were presented via Presentation software (Neurobehavioral Systems Inc., Albany CA), which allowed synchronization of scan acquisition with experiments and also recorded responses and response times (RTs). After the final scan session, participants also completed the OSIVQ (Blazhenkova and Kozhevnikov, 2009).

#### Pseudoword-shape fMRI task

We created two auditory and two visual stimuli (pseudowords and novel two-dimensional outline shapes, respectively). The auditory pseudowords were ‘lohmoh’ (rounded) and ‘keekay’ (pointed). The pseudowords were digitally recorded in a female voice using Audacity v2.0.1 (Audacity Team, 2012), with a SHURE 5115D microphone and an EMU 0202 USB external sound card, at a 44.1 kHz sampling rate. The recordings were then processed in Sound Studio (Felt Tip Inc., NY), using standard tools and default settings, edited into separate files, amplitude-normalized, and down-sampled to a 22.05 kHz sampling rate (standard for analyses of speech). Stimulus duration was 533 ms for ‘keekay’ and 600 ms for ‘lohmoh’, the differences reflecting natural pronunciation (see below). The visual stimuli were gray outline shapes on a black background (Figure 1a), each subtending approximately 1° of visual angle and presented at the center of the screen for 500ms. The selected stimuli lay near the ends of independent rounded and pointed dimensions in each modality based on empirical ratings for 537 pseudowords (McCormick et al., 2015) and 90 visual shapes (McCormick, 2015, unpublished data). These independent rating scales were converted to a single scale on which ‘most rounded’ = 1 and ‘most pointed’ = 7; ratings for the pseudowords ranged from 2.3 – 5.8 (median 4.2) and for the shapes from 1.6 – 6.7 (median 4.4). ‘Lohmoh’ was the 10^th^ most rounded pseudoword, rated 2.8, and ‘keekay’ was the 7th most pointed pseudoword, rated 5.6, and were the most similar in duration of possible pairs from the 10 pseudowords at either extreme. We used the most rounded visual shape, rated 1.6, but had to use the 11^th^ most pointed visual shape, rated 6.4, in order to match the number of protuberances (four) between the shapes. A more detailed analysis of the pseudowords and shapes can be found in Lacey et al., (2020) and the stimuli themselves are available at https://osf.io/ekpgh/. While the pseudowords differed in duration by 67ms, approximately 12%, sound segments in language naturally differ in duration and digitally altering them to match in duration can cause them to sound artificially rapid or prolonged. Although it would have been possible to match the duration of the visual shapes to whichever pseudowords they accompanied, either 533 ms or 600 ms, this would mean that shape duration would vary as a function of congruency and thus introduce a confound, particularly in the ‘attend visual’ condition (see below).

**Figure 1.**
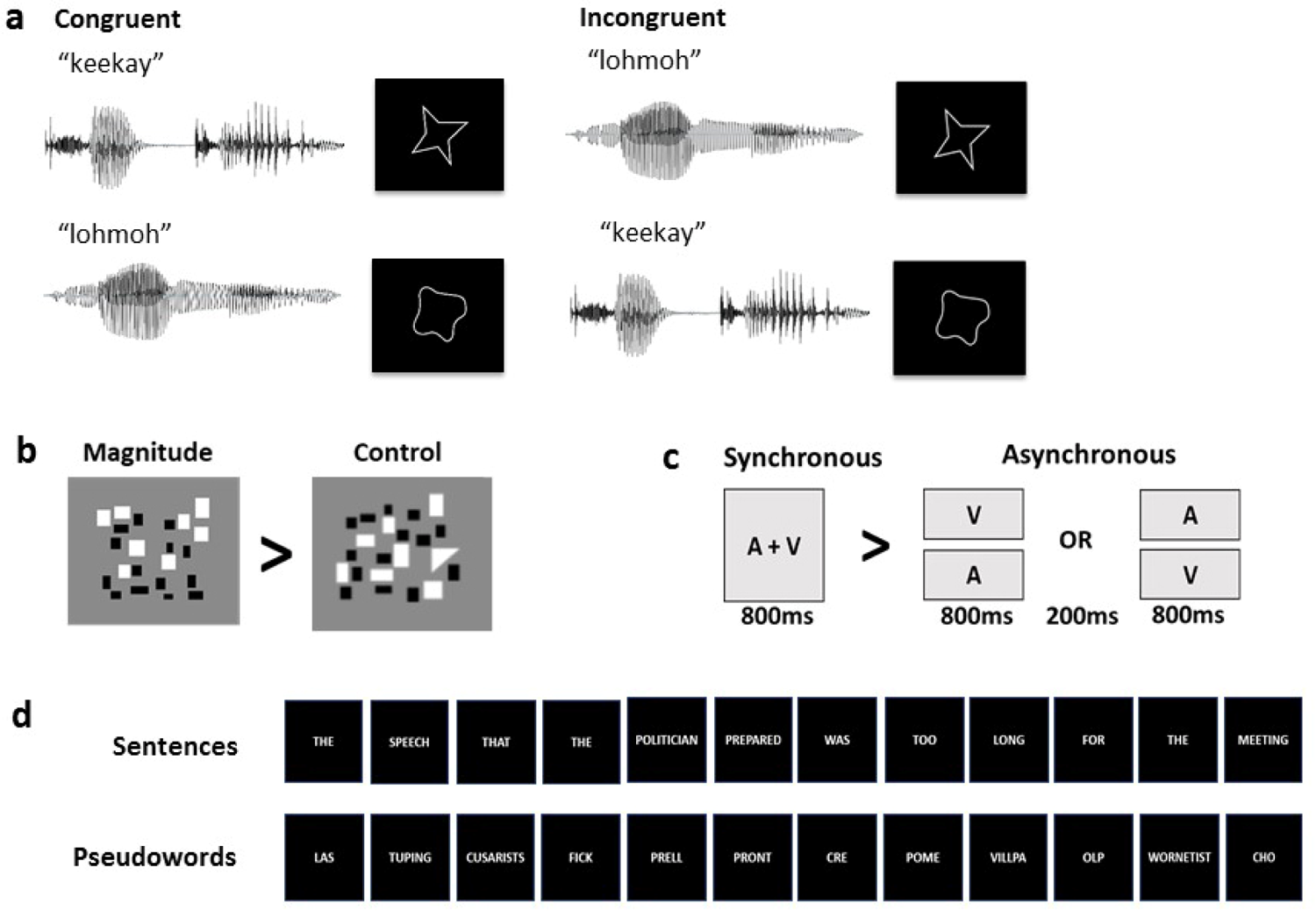
Example stimuli for (a) the sound-symbolic word-shape correspondence task and the independent localization tasks: (b) magnitude, (c) multisensory integration, (d) language.

Stimuli were presented concurrently in audiovisual pairs (Figure 1a) that were either congruent (‘keekay’/pointed shape or ‘lohmoh’/rounded shape) or incongruent (‘keekay’/rounded shape or ‘lohmoh’/pointed shape) with respect to the crossmodal pseudoword-shape (sound-symbolic) correspondence. A mirror angled over the head coil enabled participants to see the visual stimuli projected onto a screen placed in the rear magnet aperture. Auditory stimuli were presented via scanner-compatible headphones. There were four runs, each consisting of 8 task blocks (4 congruent and 4 incongruent, each block containing 10 word-shape stimuli with a 3s interval between the onset of successive stimuli), each lasting 30s and alternating with 9 rest blocks, during which a fixation cross was displayed, each lasting 16s; total run duration was 384s.

Pseudoword-shape pairs were pseudorandomly interleaved (no more than three trials in a row of the same pairing, no more than two blocks in a row of the same condition). In two of the runs, participants attended to auditory stimuli; in the other two runs, they attended to visual stimuli; the order of attended modality was counterbalanced across participants, who performed a 2AFC task in the attended modality. Participants pressed one of two buttons on a hand-held response box when they heard either ‘keekay’ or ‘lohmoh’ (attend auditory condition) or saw either the rounded or pointed shape (attend visual condition). The right index and middle fingers were used to indicate responses, counterbalanced between subjects across modalities and across rounded and pointed stimuli. Note that requiring a response on every trial and monitoring performance accuracy provides assurance that participants were attending to the stimuli. In this respect the task differs from that used by Peiffer-Smadja and Cohen (2019) in which participants only had to detect rare occurrences of visual crosses and auditory beeps rather than respond to the pseudowords and shapes. Thus, the sound-symbolic stimuli of Peiffer-Smadja and Cohen (2019) were processed incidentally and attention to these could not be guaranteed. As Peiffer-Smadja and Cohen (2019) note, the need to focus on the rare target stimuli may have modulated the processing of pseudowords and shapes, whereas in the present study, the in-scanner behavioral task involved distinguishing between stimuli in the attended modality and the effect of the unattended modality was implicit.

#### Independent localization tasks to test potential explanations of sound symbolic crossmodal correspondences

The order of these tasks was fixed, progressing from the one perceived as most difficult to the easiest: participants completed the magnitude estimation task first, then the temporal synchrony task, and finally the language task. Each of these tasks comprised two runs with a fixed stimulus order; the order of runs was counterbalanced across participants. These tasks were completed in a single session for most participants. As outlined in the Introduction, we chose these tasks to broadly reflect the three potential explanations underlying crossmodal correspondences proposed by Spence (2011): “structural”, e.g., driven by intensity or magnitude; statistical, i.e., features that regularly co-occur in the world and thus might lend themselves to multisensory integration; and semantic, i.e. driven by linguistic factors. In order to test the accounts proposed by Spence (2011), we chose localization tasks from the literature on language processing (Fedorenko et al., 2010), multisensory integration (e.g., Beauchamp, 2005a,b; van Atteveldt et al., 2007; Stevenson et al., 2010; Marchant et al., 2012; Noesselt et al., 2012; Erickson et al., 2014), and magnitude processing (Lourenco et al. 2012) as described below. A number of different task contrasts could potentially be designed to test each explanation; thus it would be difficult to test all possibilities in a single study. We made choices from the literature that seemed reasonable *a priori*, with the thinking that the results of the present study should help to refine such choices in future work (see Discussion) and with the goal of avoiding potential confounds. For example, in the multisensory task, we avoided audiovisual speech (e.g., van Atteveldt et al., 2007; Stevenson et al., 2010; Noesselt et al., 2012; Erickson et al., 2014) and familiar environmental sounds and images (e.g., Hein et al., 2007; Noppeney et al., 2008) because these would likely also involve semantic processing which we wished to test separately from multisensory integration.

Additionally, we should note that we used the framework of Spence (2011) as a starting point for a tractable set of testable hypotheses rather than an exhaustive list.

#### Magnitude estimation

In order to identify brain regions sensitive to magnitude, we used a modified form of the estimation task from Lourenco et al. (2012). In each trial of this task, participants were asked to indicate whether there were more black or white elements in a visual array of small rectangles by pressing one of two buttons on a response box. For a control task, we modified these arrays so that one item was a triangle and participants indicated whether the triangle was black or white using the same button presses as for the estimation task. There were two runs, each containing 12 active 16s-blocks (6 of each type, each block containing 4 trials lasting 1s with a 3s ISI) and alternating with 13 rest blocks, during which a fixation cross was displayed, each lasting 14s; total run duration was 374s. As can be seen in Figure 1b, the two tasks involve very similar visual displays and both required some element of visual search and the same 2AFC response, either ‘black’ or ‘white’. Thus, low-level visual, response, and motor processing were approximately equated between experimental and control tasks; the only difference was that the control task did not depend on the number of items in the array. The contrast of magnitude > control should therefore reveal areas more active during magnitude estimation than during the control task, thus identifying regions sensitive to magnitude in a domain-general magnitude system which may be relevant to assessing the magnitude along dimensions such as roundedness or pointedness. We expected that such regions would be primarily in posterior parietal cortex, particularly the IPS (Sathian et al., 1999; Eger et al., 2003; Walsh, 2003; Pinel et al., 2004; Piazza et al., 2004, 2007; Lourenco and Longo, 2012; Sokolowski et al., 2017).

#### Multisensory integration

Among a number of possible tasks that could be used to test multisensory integration, we chose one that is sensitive to the synchrony of auditory and visual stimuli, as used in many studies of audiovisual integration (e.g., Beauchamp, 2005a,b; van Atteveldt et al., 2007; Stevenson et al., 2010; Marchant et al., 2012; Noesselt et al., 2012; Erickson et al., 2014). As noted above, we attempted to avoid semantic connotations of the stimuli used in this task, such as speech or environmental sounds, with the goal of investigating non-linguistic audiovisual integration. The auditory stimulus was an 810Hz tone of 800ms duration with a 20ms on/off ramp. The visual stimulus was a gray circle (RGB values 240, 240, 240) subtending approximately 1° of visual angle and presented centrally for 800ms. In synchronous trials, auditory and visual stimuli were presented simultaneously for 800ms followed by a 3200ms blank for a total trial length of 4s, while in asynchronous trials auditory and visual stimuli were presented for 800ms each but separated by an inter-stimulus interval (ISI) of 200ms followed by an 2200ms blank, also totaling 4s (Figure 1c). During a trial, when no visual stimulus was present, the screen remained blank. Half the asynchronous trials presented the auditory stimulus first and half the visual stimulus first. There were two runs, each consisting of 12 active 16s-blocks (6 of each type, each block containing 4 trials) and alternating with 13 rest blocks each lasting 14s, during which a fixation cross was displayed; total run duration was 374s. Participants had to press a button whenever an oddball stimulus (e.g., Crottaz-Herbette and Menon, 2006), either a square or a burst of white noise, occurred; two oddballs of each type occurred in each run, one in a synchronous block and one in an asynchronous block. The contrast between synchronous and asynchronous trials was used to identify brain regions sensitive to audiovisual synchrony; we anticipated that this contrast would activate superior temporal cortex (Beauchamp, 2005a,b; van Atteveldt et al., 2007; Stevenson et al., 2010; Marchant et al., 2012; Noesselt et al., 2012; Erickson et al., 2014).

#### Language task

In this task, adapted from Fedorenko et al. (2010), we contrasted complete semantically and syntactically intact sentences with sequences of pseudowords (examples are provided in Figure 1d) to identify brain regions processing word- and sentence-level meaning (Fedorenko et al., 2010, 2011; Bedny et al., 2011). The aim of this task contrast was to reveal potential overlaps between the processing of meaning in language and the artificial pseudowords we used; for the pseudowords, meaning is implied by the correspondence of their sounds with the visual shapes, as is the case in the extensive literature on sound-symbolic crossmodal correspondences. The complete sentences and pseudoword sequences each contained 12 items, each item being presented visually for 450ms for a total stimulus duration of 5.4s. As in Fedorenko et al. (2010), participants were instructed to read each sentence/sequence and at the end of each they were visually prompted, by a cue displayed for 600ms, to press a button, following which the next trial was presented immediately. There were two runs, each consisting of 16 task blocks (8 of each type, each block containing 3 trials) of 18s duration and alternating with 17 rest blocks of 12s duration, during which a fixation cross was displayed; total run duration was 492s. We expected the contrast of complete sentences > pseudowords to reveal canonical language regions mediating both semantic and syntactic processing (see Fedorenko et al., 2010, 2011; Bedny et al., 2011), i.e., a largely left hemisphere network comprising the inferior frontal gyrus (IFG), angular gyrus (AG), and extensive sectors of the temporal lobe including the superior temporal sulcus (STS).

### Post-scan behavioral testing

As a final step, we tested whether participants reliably demonstrated the crossmodal pseudoword-shape correspondence using the implicit association test (IAT: Greenwald et al., 1998; Parise and Spence, 2012; Lacey et al., 2016). Originally devised as a test of social attitudes (Greenwald et al., 1998), the IAT has been successfully used to test the very different associations involved in crossmodal correspondences, including sound-symbolic ones (Peiffer-Smadja and Cohen, 2019; Parise and Spence, 2012; Lacey et al., 2016). The underlying principle is the same: response times (RTs) are faster if the stimuli assigned to a particular response button are congruent and slower if they are incongruent (Greenwald et al., 1998; Parise and Spence, 2012). The advantage of the IAT for testing crossmodal correspondences is that presenting each stimulus in isolation eliminates confounding by selective attention effects potentially causing slower RTs for incongruent pairings (Parise and Spence, 2012).

For the IAT, participants were presented with one of the two auditory pseudowords, ‘keekay’ or ‘lohmoh’, or one of the two visual shapes, pointed or rounded, as single items. Participants had to press one of two response buttons (the ‘left’ and ‘right’ arrows on a standard US ‘QWERTY’ keyboard) whenever they heard or saw a specific item. Thus, the key to the IAT is that there were four stimuli but only two response buttons, so that each button was used to respond to both pseudoword and shape presentations. These response button assignments could be sound-symbolically congruent (e.g., participants would press ‘left’ if they heard ‘keekay’ or saw the pointed shape, but press ‘right’ if they heard ‘lohmoh’ or saw the rounded shape) or incongruent (e.g., press ‘left’ for ‘keekay’ or the rounded shape, but press ‘right’ for ‘lohmoh’ or the pointed shape). Participants were asked to respond as quickly as possible but were expected to be slower in the incongruent condition where the pseudoword and shape assigned to each response button did not ‘go together’, i.e., were sound-symbolically unrelated. By contrast, responses were expected to be faster in the congruent condition where the pseudoword and shape assigned to each response button were sound-symbolically matched.

We used the same pseudowords and shapes as in the main fMRI experiment but the absolute size of the shapes was altered so that they subtended approximately 1° of visual angle in both the fMRI and IAT experimental set-ups. A trial consisted of a blank 1000ms followed by one of the pseudowords or one of the shapes. As before, pseudoword duration was either 533ms (‘keekay’) or 600ms (‘lohmoh’) but shapes were presented for 1000ms. A trial was terminated either by the participant pressing a response button or automatically 3500ms after stimulus onset if no response was made. On all trials, response buttons did not become active until 300ms after stimulus onset so that trials could not be terminated by rapid or accidental responses. The length of a block of trials (see next paragraph) thus varied slightly between participants but the maximum was 330s.

There were four ‘runs’, each consisting of two blocks of trials. Each block began with an instruction screen describing the task and which pseudoword and shape were assigned to each response button. This was followed by 12 practice trials (not included in the analysis) with on-screen feedback as to accuracy. Practice trials were followed by 48 active trials – with no feedback – 12 for each of the two pseudowords and two shapes, occurring in pseudorandom order. The response button assignments for all 48 trials were either congruent or incongruent. At the end of the first of the two blocks, there was a new instruction screen and the response button assignments changed from congruent to incongruent or vice-versa, depending on the first block. This was followed by a new set of 12 practice trials for the new response button assignments, and then a new set of 48 active trials. Across the four runs, two began with a congruent block followed by an incongruent block and two with the reverse, counterbalanced across participants. The pseudowords and shapes were counterbalanced across the left and right response buttons in both congruent and incongruent conditions. Across the four runs, there were 192 trials each for congruent and incongruent conditions with each pseudoword and shape occurring equally often in each condition. The IAT was presented via Presentation software which also recorded RTs, measured from stimulus onset.

Participants also completed the OSIVQ (Blazhenkova and Kozhevnikov, 2009), with the goal of examining the relationship between sound symbolism and individual differences in preferences for processing based on verbal, visual object imagery or visuospatial imagery. Such preferences have been argued to reflect fundamental styles of cognitive processing that vary between individuals and can predict abilities in many, seemingly unrelated, domains. In order to test the relationship to object or spatial imagery preference, which tend to oppose each other, we calculated the OSdiff score: the spatial score is subtracted from the object score of the OSIVQ to give a single scale on which positive/negative scores indicate a relative preference for object/spatial imagery respectively (Lacey et al., 2011, 2014, 2017; Occelli et al., 2014). The relationship to verbal preference was tested using the verbal score from the OSIVQ.

### Image acquisition

MR scans were performed on a 3 Tesla Siemens Trio TIM whole body scanner (Siemens Medical Solutions, Malvern, PA), using a 12-channel head coil. T2*-weighted functional images were acquired using a single-shot, gradient-recalled, echoplanar imaging (EPI) sequence for blood oxygenation level-dependent (BOLD) contrast. For all functional scans, 34 axial slices of 3.1mm thickness were acquired using the following parameters: repetition time (TR) 2000ms, echo time (TE) 30ms, field of view (FOV) 200mm, flip angle (FA) 90°, in-plane resolution 3.125×3.125mm, and in-plane matrix 64×64. High-resolution 3D anatomic images were acquired using an MPRAGE sequence (TR 2300ms, TE 3.02ms, inversion time 1100ms, FA 8°) comprising 176 sagittal slices of 1mm thickness (FOV 256mm, in-plane resolution 1×1mm, in-plane matrix 256×256). Once magnetic stabilization was achieved in each run, the scanner triggered the computer running Presentation software so that the sequence of experimental trials was synchronized with scan acquisition.

### Image processing and analysis

Image processing and analysis was performed using BrainVoyager QX v2.8.4 (Brain Innovation, Maastricht, Netherlands). Each participant’s functional runs were real-time motion-corrected utilizing Siemens 3D-PACE (prospective acquisition motion correction). Functional images were preprocessed employing cubic spline interpolation for slice scan time correction, trilinear-sinc interpolation for intra-session alignment of functional volumes, and high-pass temporal filtering to 2 cycles per run to remove slow drifts in the data without compromising task-related effects. Anatomic 3D images were processed, co-registered with the functional data, and transformed into Talairach space (Talairach and Tournoux, 1988). Talairach-normalized anatomic data sets from multiple scan sessions were averaged for each individual, to minimize noise and maximize spatial resolution.

For group analyses, the Talairach-transformed data were spatially smoothed with an isotropic Gaussian kernel (full-width half-maximum 4mm). The 4mm filter is within the 3-6mm range recommended to reduce the possibility of blurring together activations that are in fact anatomically and/or functionally distinct (White et al., 2001), and the ratio of the smoothing kernel to the spatial resolution of the functional images (1.33) matches that of studies in which larger smoothing kernels were used (Mikl et al., 2008). Runs were percent signal change normalized (i.e., the mean signal value for each voxel’s time course was transformed to a value of 100, so that the individual values fluctuated around that mean as percent signal deviations).

For display of group activations, we created a group average brain. We first selected a representative (target) Talairach-normalized brain from the 19-participant group. We then individually aligned the 18 remaining participants’ Talairach-normalized brains to this target (co-registration to match the gyral and sulcal pattern, followed by sinc interpolation). These 18 aligned brains were then averaged. This 18-subject average brain was then averaged with the target brain, creating a single Talairach template, with 1mm isotropic resolution, which was used to display the activations for the 19-subject group. This group average brain was displayed using the real-time volume rendering option in BrainVoyager QX. For statistical analysis, the 19-subject Talairach template was manually segmented in order to create a group average cortical ‘mask’ file with 3mm spatial resolution, equivalent to the spatial resolution of the functional data files.

Statistical analyses of group data used general linear models (GLMs) treating participant as a random factor (so that the degrees of freedom equal n-1, i.e. 18), followed by pairwise contrasts. This analysis allows generalization to untested individuals. Correction for multiple comparisons within a cortical mask (corrected p < .05) was achieved by imposing a threshold for the volume of clusters comprising contiguous voxels that passed a voxel-wise threshold of p < 0.001, using a 3D extension (implemented in BrainVoyager QX) of the 2D Monte Carlo simulation procedure described by Forman et al. (1995). This stringent voxel-wise threshold is recommended to avoid potential problems of false positives and also permits more accurate spatial localization of activation clusters than is possible with more lenient thresholds (Woo et al., 2014; Eklund et al., 2016). Activations were localized with respect to 3D cortical anatomy with the help of an MRI atlas (Duvernoy, 1999).

In reporting activations, we do not provide ‘hotspot’ or ‘t_max_’ coordinates (i.e. the voxel with the largest t-value) because the statistical significance of specific voxels is not tested against other voxels within the activation (Woo et al., 2014). Instead, we provide the ‘center of gravity’ (CoG) coordinates since these orient the reader to the anatomical location but are statistically neutral.

Where an activation spans several anatomical locations, we provide an extended description (see Table 1). Likewise, in order to compare the present results to previous studies, we have not computed the Euclidean distance between sets of coordinates since, in most cases, this would involve comparing t_max_ to CoG and therefore we would not be comparing like to like. Instead, we plotted the coordinates reported in previous studies onto our data in order to assess whether these fell within the activations we found here, such that we could reasonably say that there was some degree of overlap in the activations.

**Table 1.**
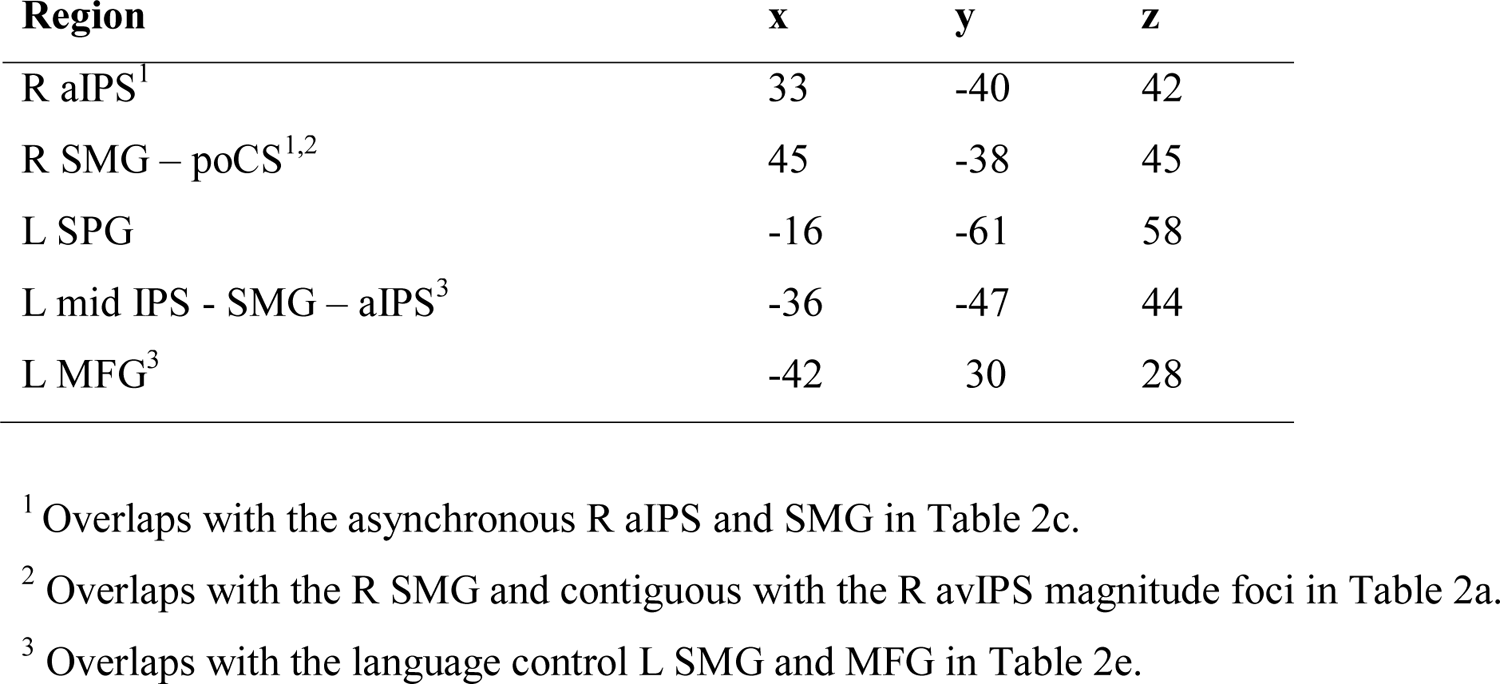
Incongruency effects: pseudoword-shape incongruency-related activations in the ‘attend auditory’ condition within the cortical mask (voxel-wise threshold p < 0.001, cluster-corrected p < 0.05 cluster threshold 5 voxels); x,y,z, Talairach coordinates for centers of gravity.

| **Region** | **x** | **y** | **z** |
| --- | --- | --- | --- |
| R aIPS<sup>1</sup> | 33 | -40 | 42 |
| R SMG – poCS<sup>1,2</sup> | 45 | -38 | 45 |
| L SPG | -16 | -61 | 58 |
| L mid IPS - SMG – aIPS<sup>3</sup> | -36 | -47 | 44 |
| L MFG<sup>3</sup> | -42 | 30 | 28 |

<sup>1</sup> Overlaps with the asynchronous R aIPS and SMG in Table 2c.
<sup>2</sup> Overlaps with the R SMG and contiguous with the R avIPS magnitude foci in Table 2a.
<sup>3</sup> Overlaps with the language control L SMG and MFG in Table 2e.

## RESULTS

### Behavioral

#### In-scanner tasks

##### Pseudoword-shape task

Two-way (congruency, attended modality) repeated-measures ANOVA (RM-ANOVA) showed that mean (±sem) accuracy was not significantly different between the ‘attend auditory’ (97.4±0.7%) and ‘attend visual’ conditions (97.6±0.6%: F_1, 18_ = .05, p = .8), nor between congruent (97.7±0.5%) and incongruent (97.3±0.6%) trials (F_1, 18_ = .9, p = .3). There was a significant interaction between the attended modality and congruency (F_1,18_ = 9.6, p = 0.006); this was due to greater accuracy for congruent compared to incongruent trials in the ‘attend auditory’ condition (98.2% vs 96.5%; t_18_ = 2.6, p = 0.018) but not the ‘attend visual’ condition (97.1% vs 98%; t_18_ = −1.9 p = 0.06: note that we are only concerned with these two comparisons, thus Bonferroni-corrected alpha = 0.025). We also performed non-parametric tests on the accuracy data since the ‘ceiling’ effects in all conditions indicate that the data were likely not normally distributed: these confirmed the result of the RM-ANOVA.

Analysis of RTs excluded trials for which there was no response (.6% of all trials) or an incorrect response (2.5% of responses), and further excluded trials for which the RT was more than ± 2.5 standard deviations from the individual participant mean (2.6% of correct response trials). RM-ANOVA showed that RTs were faster for the ‘attend visual’ (474±21ms) compared to the ‘attend auditory’ condition (527±23ms: F_1,18_ = 21.3, p < 0.001) and for congruent (489±21ms) compared to incongruent trials (513±22ms: F_1,18_ = 18.7, p < 0.001). There was a significant interaction between the attended modality and congruency (F_1,18_ = 9.2, p < 0.007) in which both auditory (545ms) and visual (480ms) incongruent RTs were slower than congruent RTs in the same modality (auditory 509ms, t_18_ = −4.0, p = 0.001; visual 469ms, t_18_, p = 0.009) but the absolute difference was greater for auditory than visual RTs (36ms vs 11ms).

Note that, despite the highly repetitive nature of the pseudoword and shape stimuli, it is unlikely that participants stopped paying attention to them given the high accuracy rates (> 97% in all conditions) and the fact that responses were made on 99.4% of trials. In addition, the mean auditory RTs (527ms overall, and 509ms vs. 545ms for congruent vs incongruent trials) suggest that participants often responded before the auditory stimulus was complete (duration = 533ms vs 600ms for ‘keekay’ vs. ‘lohmoh’). This was likely unavoidable given that participants only had to identify which of the two, highly repeated, pseudowords they heard; but the first syllable of each pseudoword, either ‘kee-’ or ‘loh-’, also respects the class of the entire word, either pointed or rounded respectively. Thus, we do not believe that the results are affected by such rapid responses since our analyses reveal that while being scanned, participants still exhibited significant behavioral congruency effects related to sound symbolism, more prominently in the attend-auditory than the attend-visual condition. On a related note, we do not believe that the longer auditory RTs indicate that the pseudoword detection task was more difficult than the visual shape task. The auditory task was to press one button whenever participants heard ‘keekay’ and the other for ‘lohmoh’, i.e., it was not an inherently difficult task and, as we point out above, the auditory RT results suggest that participants often responded before the end of the pseudoword which, given the high accuracy level (equal to that in the visual shape task), suggests that the task was not difficult. The longer auditory RTs appear to be driven by the incongruent trials rather than inherent task difficulty.

##### Independent localization tasks

For the magnitude task, there was no significant difference in accuracy between the magnitude estimation (92.2±2.0%) and control (96.4±1.3%) tasks (t_18_ = −1.8, p = 0.1) although RTs were significantly faster for the control task (900±45ms) compared to the magnitude task (991±53ms; t_18_ = 3.4, p < 0.01). Because of the low number of oddball trials (four for each participant) in the multisensory task, we conducted a non-parametric Wilcoxon test, which showed no significant difference between detection of synchronous (90.1±3.9%) and asynchronous (85.5±5.5%) oddballs (Z = −1.4, p = 0.2). In the language task, participants were equally accurate in responding to the visual cue at the end of each sentence (mean ± sem: 98.4±1.0%) or pseudoword string (97.7±1.1%; t18 = 0.9, p = 0.4). These analyses allow us to infer that participants paid attention to the stimuli used in the independent localization tasks.

### Post-scan pseudoword-shape IAT

An RM-ANOVA with factors of modality (auditory pseudowords, visual shapes) and response key association (congruent, incongruent) showed that accuracy was higher for the auditory pseudowords (95.8±0.8%) compared to the visual shapes (91.9±0.8%: F_1,18_ = 31.8, p < 0.001) and when response key associations were congruent (95.3±0.7%) compared to incongruent (92.4±1.2%: F_1,18_ = 5.7, p = 0.03). The modality x response key association interaction was not significant (F_1,18_ < .01, p = 0.9).

Analysis of RTs excluded trials for which there was no response or which failed to log (.7% of all trials), or incorrect responses (6.2% of responses), and further excluded trials for which the RT was more than ± 2.5 standard deviations from the individual participant mean (2.7% of correct response trials). RM-ANOVA showed that RTs were faster for the visual (606±19ms) compared to the auditory (702±21ms) stimuli (F_1,18_ = 31.15, p < 0.001) and when the response key associations were congruent (580±18ms) compared to incongruent (728±22ms: F_1,18_ = 64.5, p < 0.001). The modality x response key association interaction was not significant (F_1,18_ = 0.5, p = 0.5).

These analyses provide converging evidence, using a different approach from that employed during scanning, that participants experienced sound-symbolic congruency effects with the stimuli that were used.

### Imaging

#### Pseudoword-shape task

To test for regions involved in processing the sound-symbolic word-shape correspondence, we tested for voxels showing differential activation for congruent and incongruent trials (C > I or I > C), i.e., congruency or incongruency effects, respectively. These analyses were performed independently for the attend-auditory and attend-visual runs. Within the group average cortical mask, at a voxel-wise threshold of p < 0.001, there were no activations that survived correction for multiple comparisons in the ‘attend visual’ condition, either for the C > I contrast or its reverse, the I > C contrast.

However, in the ‘attend auditory’ condition, while there was no effect favoring the congruent over the incongruent condition, several regions showed greater activation for incongruent, compared to congruent, trials (the I > C contrast) within the cortical mask (voxel-wise threshold p < 0.001, cluster-corrected p < 0.05, cluster threshold 5 voxels). These regions included bilateral foci in the anterior IPS (aIPS) extending on the left into the mid-IPS and the SMG, and activations in the right SMG extending into the postcentral sulcus (poCS), in the left SPG and the left MFG (Table 1; Figure 2). Representative time-course curves for the I > C contrast in these regions are displayed in Figure 6A. These show greater activation for incongruent compared to congruent trials in all regions except for the left SPG region in which there was differential *de*activation. Since it is not clear how to interpret differential deactivations, we do not consider this SPG focus further^2^. The neural incongruency effects were associated with a behavioral congruency effect in the ‘attend auditory’ condition in which responses to congruent trials were faster and more accurate than those to incongruent trials; this is consistent with the processing of incongruent trials being more effortful. The absence of a neural (in)congruency effect in the ‘attend visual’ condition is consonant with the absence of a congruency effect for accuracy data in this condition; although there was a congruency effect for RTs, this effect was significantly smaller in the ‘attend visual’ condition compared to the ‘attend auditory’ condition (see above).

**Figure 2.**
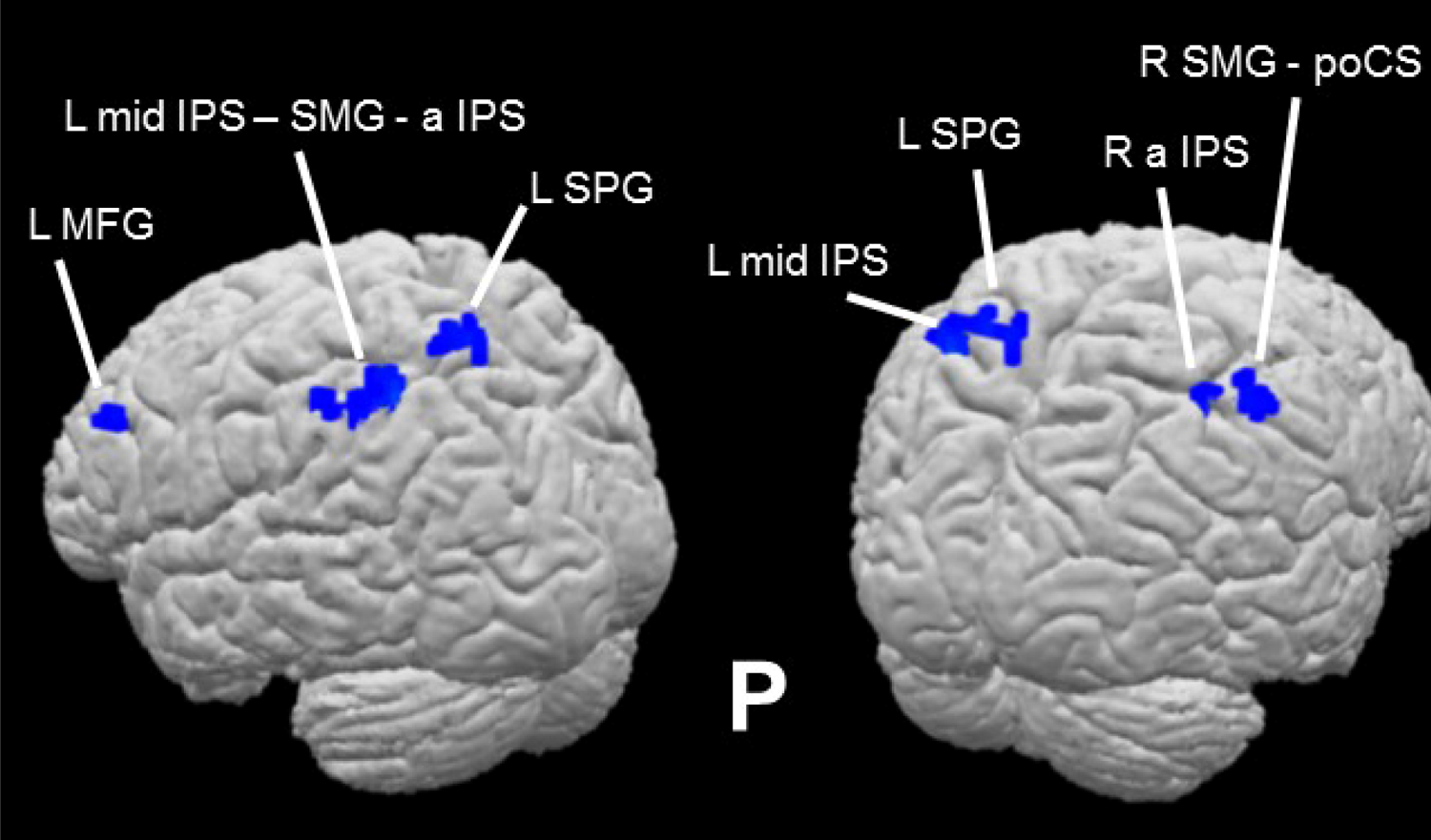
Sound-symbolic incongruency effects: pseudoword-shape incongruency-related activations in the ‘attend auditory’ condition (Table 1).

### Independent localization tasks

#### Magnitude estimation

The contrast of magnitude estimation > control within the cortical mask (voxel-wise threshold p < 0.001, cluster-corrected p < 0.05, cluster threshold 7 voxels) showed exclusively right hemisphere activity in the SMG, the SPG extending into the IPS, and the middle occipital gyrus (MOG) extending through the intra-occipital sulcus (IOS) to the superior occipital gyrus (SOG: Table 2a; Figure 3). These loci are consistent with activations reported in previous studies of magnitude processing (Eger et al., 2003; Pinel et al., 2004; Piazza et al., 2004, 2007) and a recent meta-analysis (Sokolowski et al., 2017). The right MOG region is close to foci previously implicated in subitizing (Sathian et al., 1999) or that showed adaptation to magnitude (Piazza et al., 2007) while the right SPG-IPS region is close to a region involved in counting visual objects (Sathian et al., 1999).

**Figure 3.**
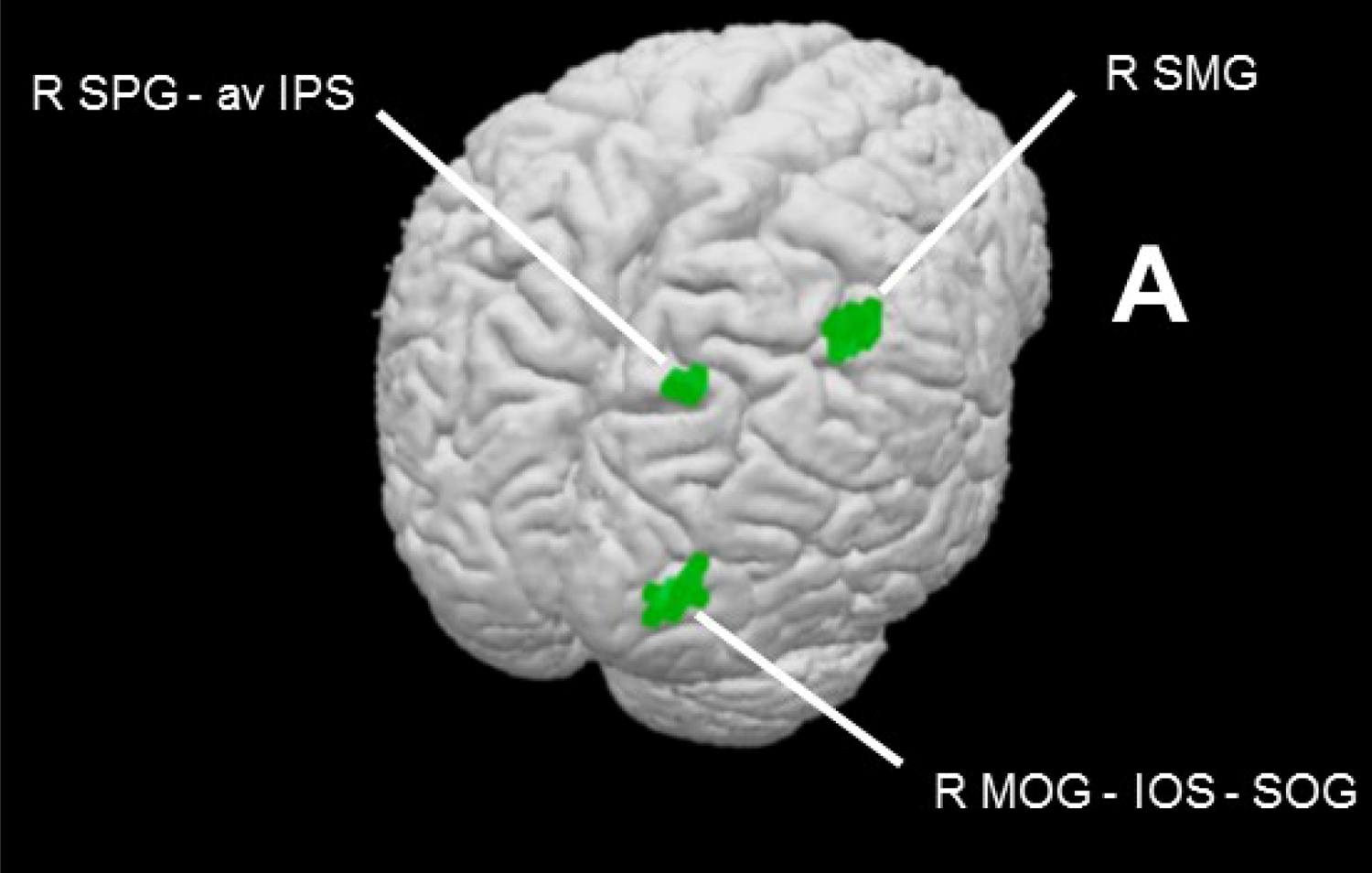
Magnitude task effects within cortical mask (voxel-wise threshold p < 0.001, cluster-corrected p < 0.05, cluster threshold 7 voxels). Contrast of magnitude estimation > control reveals magnitude network (Table 2a).

**Table 2.**
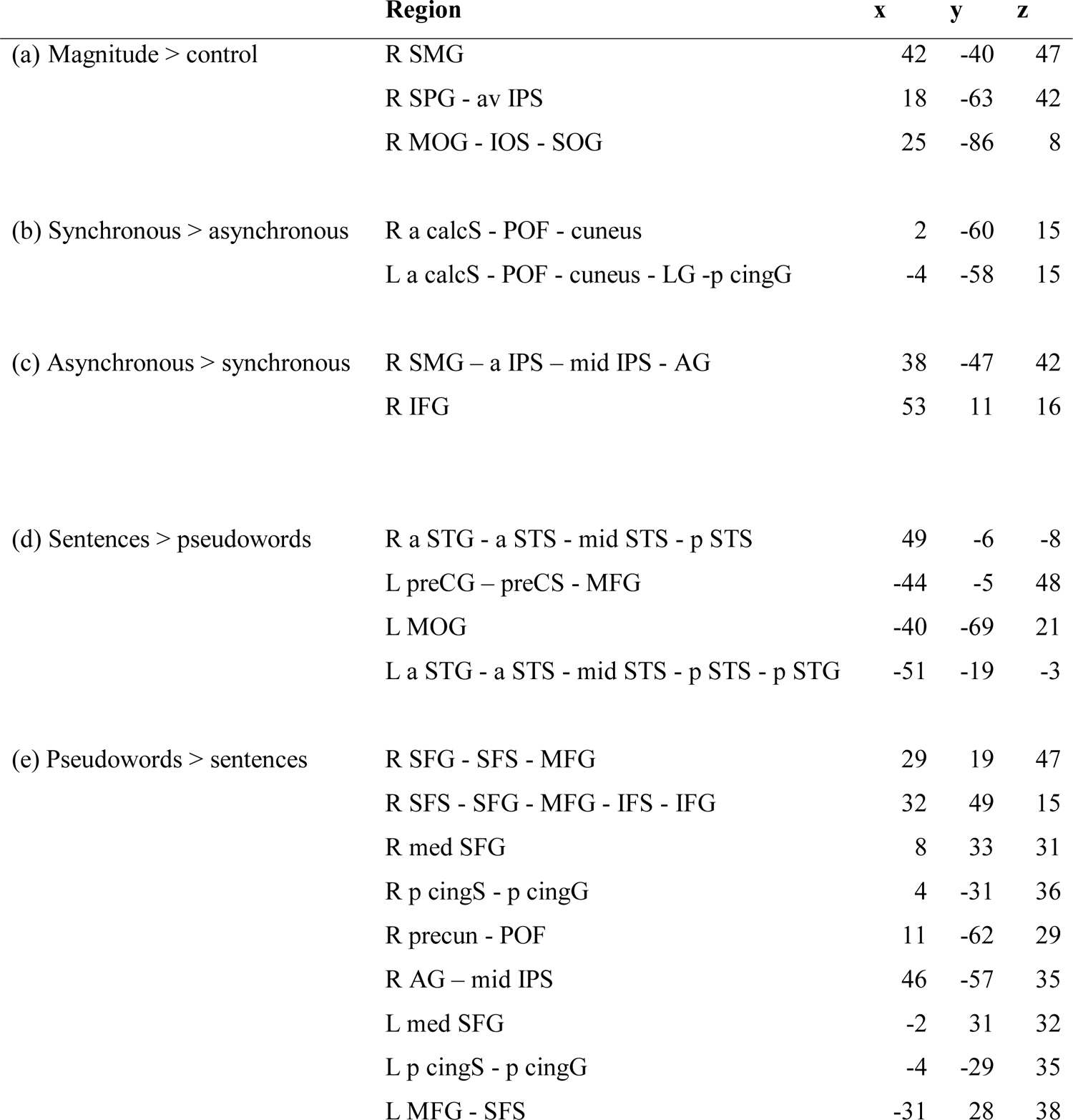
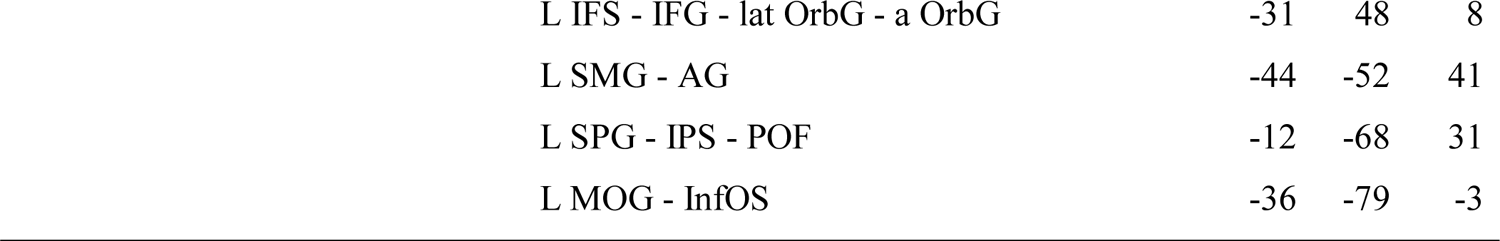
Activations on independent localization tasks: magnitude (a), multisensory integration (b,c), and language (d,e); all within a cortical mask, voxel-wise threshold p < 0.001, cluster-corrected p < 0.05, cluster thresholds = magnitude, 7 voxels; multisensory integration, 8 voxels; language, 9 voxels; x,y,z Talairach coordinates for centers of gravity.

|  | **Region** | **x** | **y** | **z** |
| --- | --- | --- | --- | --- |
| (a) Magnitude > control | R SMG | 42 | -40 | 47 |
|  | R SPG - av IPS | 18 | -63 | 42 |
|  | R MOG - IOS - SOG | 25 | -86 | 8 |
| (b) Synchronous > asynchronous | R a calcS - POF - cuneus | 2 | -60 | 15 |
|  | L a calcS - POF - cuneus - LG -p cingG | -4 | -58 | 15 |
| (c) Asynchronous > synchronous | R SMG – a IPS – mid IPS - AG | 38 | -47 | 42 |
|  | R IFG | 53 | 11 | 16 |
| (d) Sentences > pseudowords | R a STG - a STS - mid STS - p STS | 49 | -6 | -8 |
|  | L preCG – preCS - MFG | -44 | -5 | 48 |
|  | L MOG | -40 | -69 | 21 |
|  | L a STG - a STS - mid STS - p STS - p STG | -51 | -19 | -3 |
| (e) Pseudowords > sentences | R SFG - SFS - MFG | 29 | 19 | 47 |
|  | R SFS - SFG - MFG - IFS - IFG | 32 | 49 | 15 |
|  | R med SFG | 8 | 33 | 31 |
|  | R p cingS - p cingG | 4 | -31 | 36 |
|  | R precun - POF | 11 | -62 | 29 |
|  | R AG – mid IPS | 46 | -57 | 35 |
|  | L med SFG | -2 | 31 | 32 |
|  | L p cingS - p cingG | -4 | -29 | 35 |
|  | L MFG - SFS | -31 | 28 | 38 |

|  |  |  |  |
| --- | --- | --- | --- |
| L IFS - IFG - lat OrbG - a OrbG | -31 | 48 | 8 |
| L SMG - AG | -44 | -52 | 41 |
| L SPG - IPS - POF | -12 | -68 | 31 |
| L MOG - InfOS | -36 | -79 | -3 |

#### Multisensory synchrony

The contrast of synchronous > asynchronous within the cortical mask (voxel-wise threshold p < 0.001, cluster-corrected p < 0.05, cluster threshold 8 voxels) revealed bilateral activations in the anterior calcarine sulcus extending through the POF to the cuneus; in the left hemisphere, this activation further encompassed foci in the lingual and posterior cingulate gyri (Table 2b; Figure 4). The reverse contrast, asynchronous > synchronous, resulted in two right hemisphere activations, one in inferior parietal cortex extending across the SMG and AG and into the IPS, and one in the IFG (Table 2c; Figure 4). While some previous studies have shown greater activation for synchronous compared to asynchronous audiovisual stimuli, others have shown the reverse, and in some cases preference for synchrony and asynchrony occurred at foci in proximity to each other (e.g. Stevenson et al., 2010). The regions most consistently reported as being sensitive to synchrony in previous studies were in the STS and/or STG – we did not observe activations in these regions on the multisensory synchrony task here, perhaps reflecting differences in stimuli and tasks (see Discussion). However, the right IFG area that was more active for asynchronous than synchronous stimuli in the present study was close to a region identified on the meta-analysis of Erickson et al. (2014) as exhibiting a preference for audiovisual stimuli characterized by either content incongruency or asynchrony.

**Figure 4.**
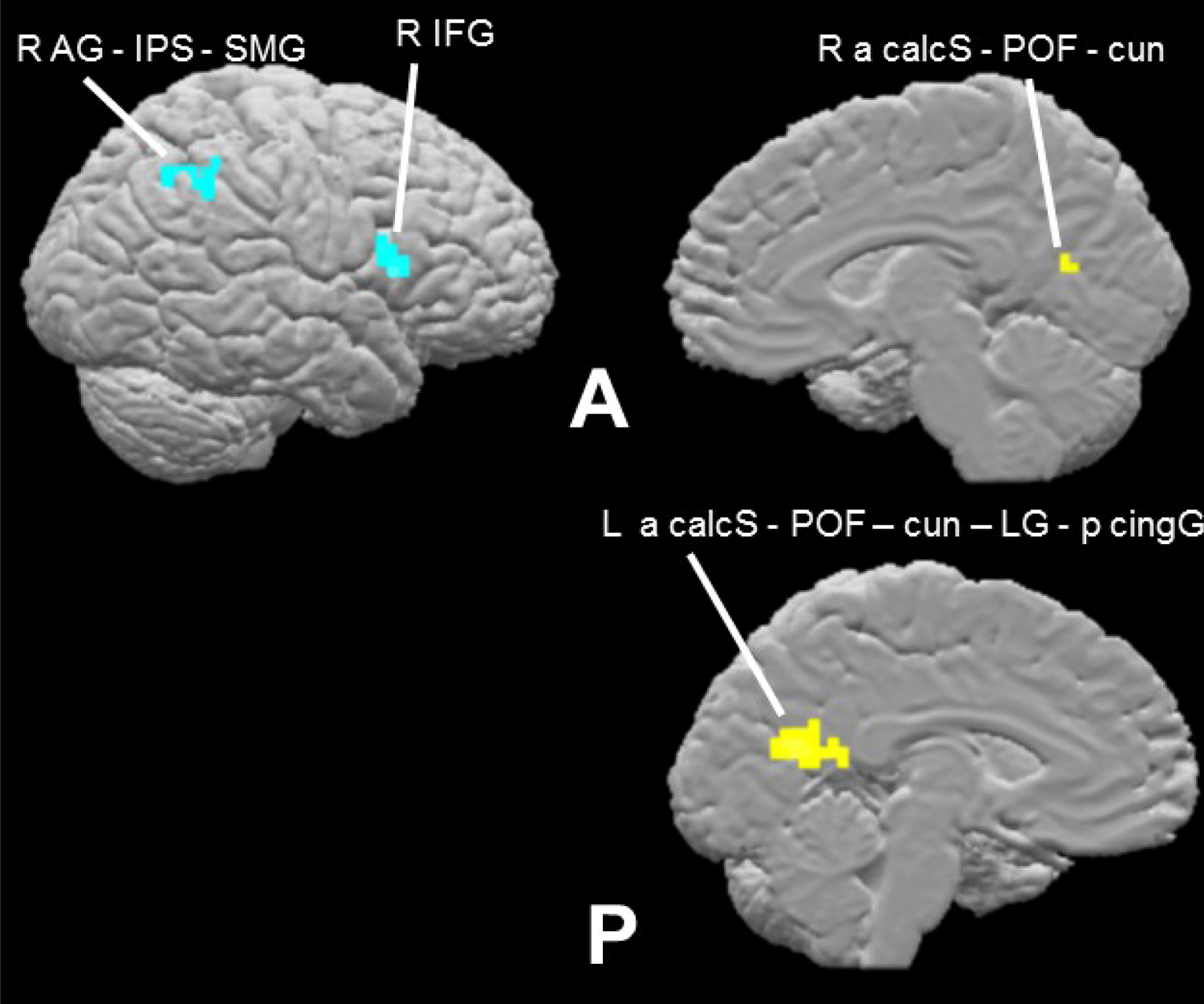
Multisensory synchrony task effects within cortical mask (voxel-wise threshold p < 0.001, cluster-corrected p < 0.05, cluster threshold 8 voxels). Contrast of synchronous > asynchronous (turquoise) reveals putative integration network (Table 2b); asynchronous > synchronous (yellow: Table 2c).

#### Language

As expected, the contrast of complete sentences > pseudowords within the group average cortical mask described in the Methods section (voxel-wise threshold p < 0.001, cluster-corrected p < 0.05, cluster threshold 9 voxels) revealed large activations bilaterally along the STS, extending into parts of the superior temporal gyrus (STG); this activation extended more posteriorly on the left than the right. Additional activations were noted on the left precentrally and in the middle occipital gyrus (Table 2d; Figure 5). Thus, the language task broadly replicated the previously reported language network (Fedorenko et al., 2010, 2011).

**Figure 5.**
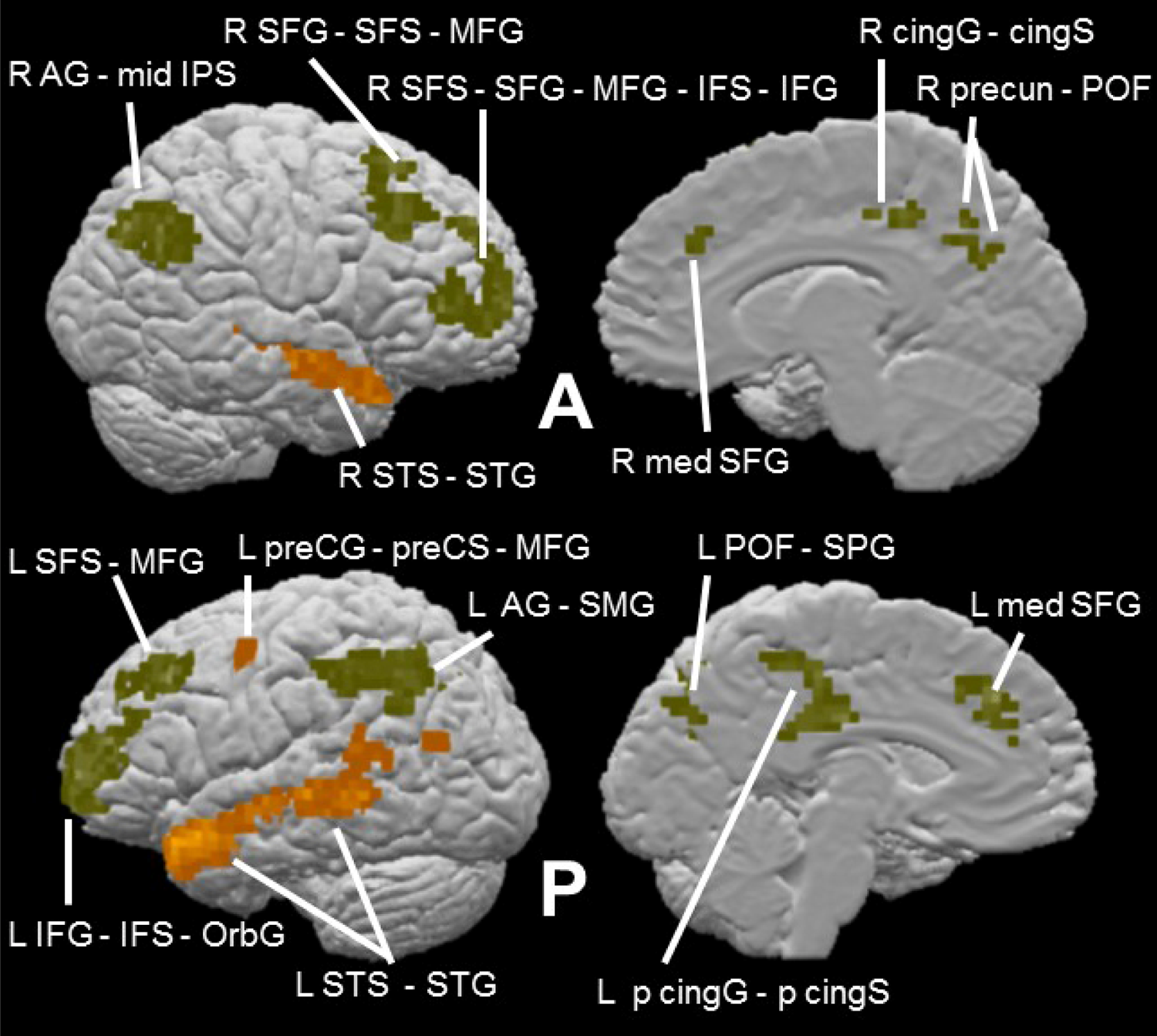
Language task effects within cortical mask (voxel-wise threshold p < 0.001, cluster-corrected p < 0.05, cluster threshold 9 voxels). Contrast of sentences > pseudowords (orange) reveals semantic network (Table 2d); pseudowords > sentences (olive) (Table 2e).

Although not part of the original design of the present study, the contrast of pseudowords > sentences may reflect the greater role of phonological processing in reading strings of pseudowords compared to normal intact sentences (Fedorenko et al., 2010). Given that our auditory stimuli were pseudowords, we were interested in also examining this reverse contrast. The contrast of pseudowords > sentences within the cortical mask (voxel-wise threshold p < 0.001, cluster-corrected p < 0.05, cluster threshold 9 voxels) revealed large, bilateral frontoparietal activations: in and around the superior frontal sulcus (SFS) extending all the way into the IFG, and in the AG extending into the supramarginal gyrus (SMG) on the left and the IPS on the right. Smaller activations were also found along the medial surface of the left hemisphere in frontal and posterior cingulate cortex and in the parieto-occipital fissure (POF) (Table 2e; Figure 5). Consistent with a phonological basis, the left IFG activation on this contrast is close to an IFG focus showing greater activity for visually presented pseudowords relative to concrete words (Binder et al., 2005), a contrast similar to that used here. Additionally, the left SMG activation on this contrast is at a site reported to be involved in phonological processing for visually presented words (Price et al., 1997; Wilson et al., 2011). However, reading pseudowords is also more effortful than reading complete sentences, perhaps due to mapping unfamiliar orthographic representations to phonological representations. Therefore, another possibility is that some or all of the activations on this contrast might reflect this additional effort, particularly since many of the activations (principally those in the frontal cortex bilaterally) were also found in regions identified as part of the frontoparietal domain-general, ‘multiple demand’ system (Duncan, 2013; Fedorenko et al., 2013). We return to this point in the Discussion.

### Overlap of incongruency effect with independent localization task contrasts

Our approach to distinguishing between the competing potential explanatory accounts examined here was to look for overlap between areas showing pseudoword-shape (in)congruency effects and areas revealed by the independent language, magnitude, and multisensory tasks. Note that we assessed such overlap at voxel-wise thresholds of p < 0.001 for the activations being compared. Overlaps at this strict threshold, avoiding potential false positives and allowing more accurate spatial localization than at more liberal thresholds (Woo et al., 2014; Eklund et al., 2016), would provide support for candidate explanations. However, the presence of an overlap does not guarantee that the same process underlies both tasks at that locus nor that the same neuronal populations are activated; moreover, note that absence of overlaps would not allow such accounts to be definitively ruled *out*. The evidence of overlaps should be regarded as indicative and supportive of further research into the functional roles of those loci.

In the magnitude task, the main magnitude estimation > control contrast overlapped with the right SMG incongruency region and was also contiguous with the right aIPS incongruency region (see notes to Table 1; Figure 7). The BOLD signal time-course for the right SMG region showed similar response patterns for the sound-symbolic and magnitude tasks: greater activation for incongruent and magnitude estimation trials, respectively, relative to the corresponding comparison conditions (Figure 6A and B).

**Figure 6.**
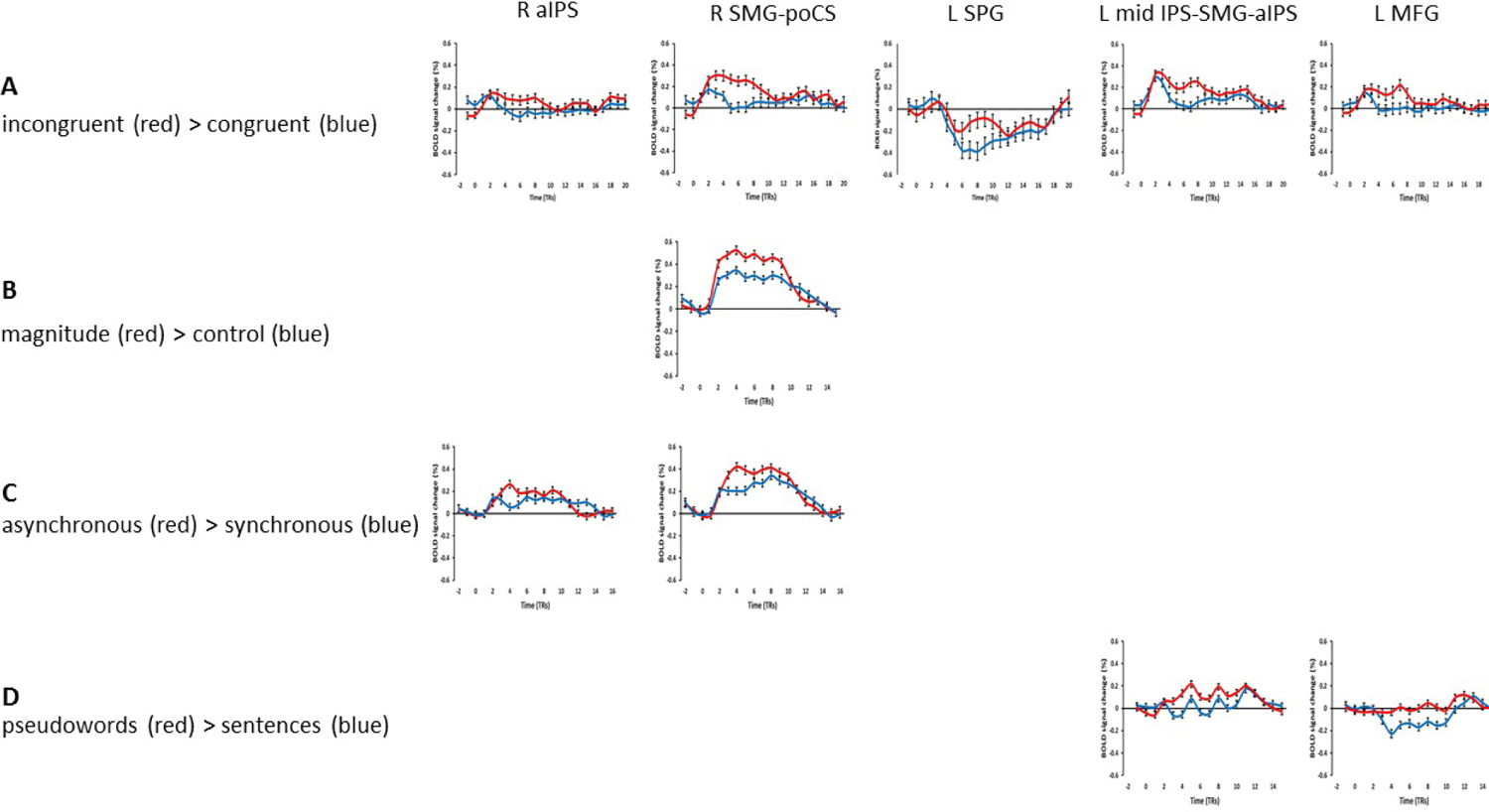
Representative time-course curves for regions showing sound-symbolic incongruency effects in (A) the incongruent > congruent contrast and (B-D) contrasts from the independent functional contrasts with which they shared an overlap zone.

In the multisensory task, the contrast of asynchronous > synchronous showed overlaps with the right aIPS and SMG incongruency regions (see notes to Table 1; Figure 7). BOLD signal time-courses for these regions showed that both exhibited similar response patterns for the sound-symbolic and multisensory synchrony tasks: greater activation for incongruent and asynchronous trials, respectively, relative to the corresponding comparison conditions (Figure 6A and C). Since the multisensory task did not reveal the STS activity that was expected on the basis of previous studies (Beauchamp, 2005a,b; van Atteveldt et al., 2007; Stevenson et al., 2010; Marchant et al., 2012; Noesselt et al., 2012; Erickson et al., 2014), we also carried out an ROI analysis in bilateral STS ROIs, chosen because they are sensitive to audiovisual integration of non-speech stimuli (tools and musical instruments and their associated sounds) and also behaviorally relevant in that activation profiles predict task performance (Werner and Noppeney, 2010).

**Figure 7.**
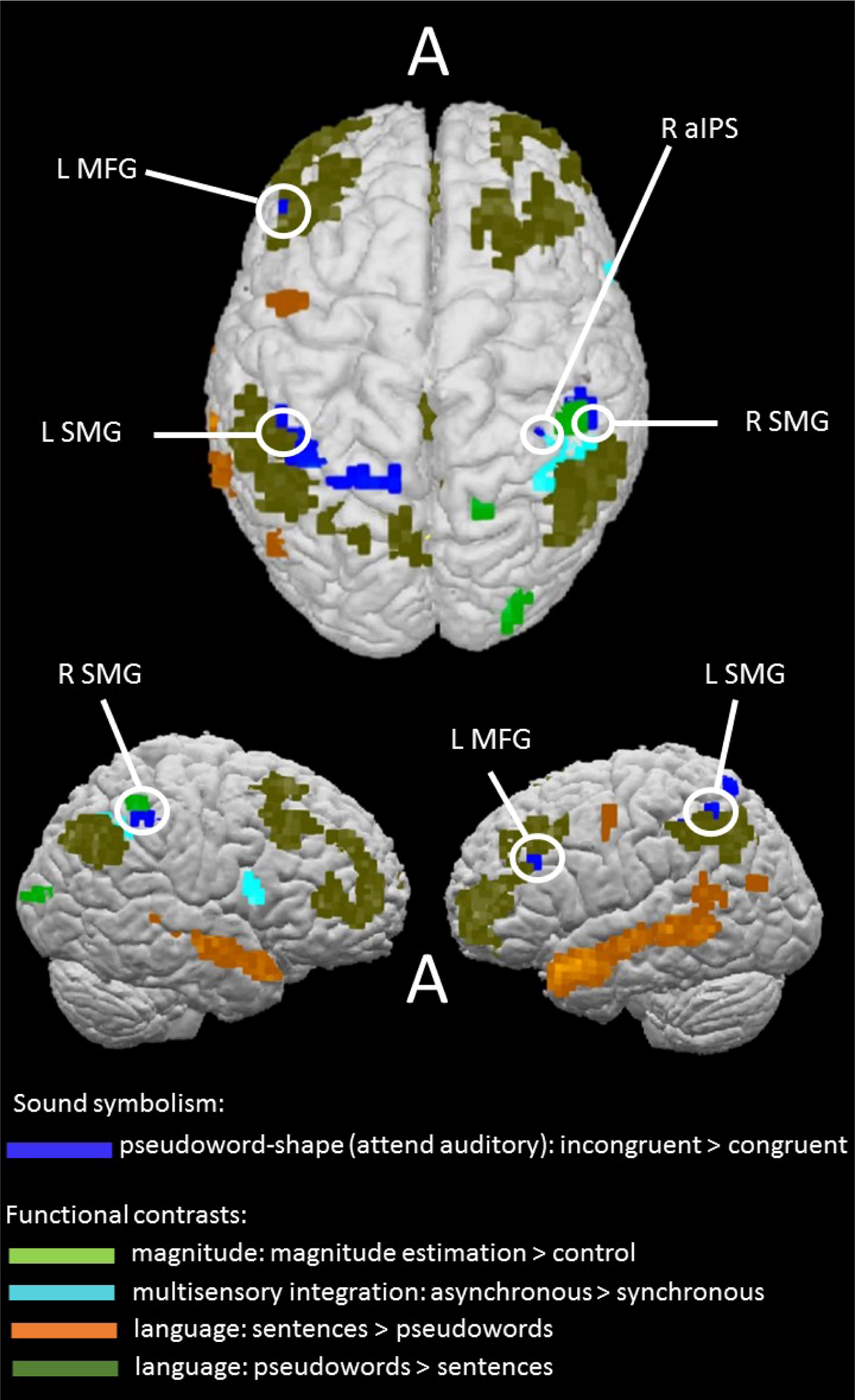
Sound-symbolic incongruency effects (blue) in relation to independent functional contrasts for magnitude processing (green), multisensory integration (turquoise), and language: sentences > pseudowords (orange), pseudowords > sentences (olive). Circles indicate areas of overlap or contiguity between incongruency effects and independent functional contrasts (see notes to Table 1).

These ROIs, which were also used in a prior study from our group on the pitch-elevation crossmodal correspondence (McCormick et al., 2018), comprised cubes of 15mm side, centered on coordinates from Werner and Noppeney (2010; Talairach coordinates −57, −41, 14; 52, −33, 9; MNI coordinates were transformed into Talairach space using the online tool provided by Lacadie et al., 2008). However, the contrast of incongruent > congruent was not significant in these ROIs, either in the ‘attend auditory’ condition (left: t_18_ = 0.3, p = 0.8; right: t_18_ = 0.4, p = 0.7) or the ‘attend visual’ condition (left: t_18_ = 1.6, p = 0.1; right: t_18_ = 1.2, p = 0.2).

Finally, none of the regions showing sensitivity to the pseudoword-shape correspondence overlapped or were contiguous with (i.e. shared a common edge or vertex) any area revealed by the language contrasts of sentences > pseudowords or the multisensory contrast of synchronous > asynchronous (Figure 7). Instead, regions showing pseudoword-shape incongruency effects showed overlaps and contiguities with the control conditions from the language and multisensory tasks, i.e. when the primary contrasts for these conditions were reversed (see notes to Table 1; Figure 7). The contrast of pseudowords > complete sentences from the language task revealed overlap with incongruency effects in the left MFG and SMG. Comparing the BOLD signal time-courses for these regions (Figure 6A and D) suggests that the left SMG region may be the more relevant of the two since it showed greater activation on the reverse contrasts. By contrast, the left MFG showed different response patterns in the sound-symbolic and language tasks: while there was greater activation for incongruent than congruent trials in the former task, it barely responded to pseudowords while deactivating to the sentence trials in the language task.

Although the pseudoword condition likely involves processing the phonological form of the pseudowords rather than the semantic or syntactic aspects of natural language (Fedorenko et al., 2010), the pseudowords > sentences contrast may have limitations as a formal test for phonological processing (see Discussion). Therefore, we also conducted a region-of-interest (ROI) analysis in which we created bilateral SMG ROIs, chosen because they were shown to be functionally involved in phonological processing using transcranial magnetic stimulation (TMS; Hartwigsen et al., 2010). The ROIs consisted of cubes of 15mm side, centered on coordinates from Hartwigsen et al. (2010; Talairach coordinates −45, −37, 42; 45, −36, 42; MNI coordinates were transformed into Talairach space as above). Although the SMG was part of the inferior parietal cluster of sound-symbolic incongruency-related activations in both hemispheres, the ROI approach allows a more specific test of the relationship to phonology. The contrast of incongruent > congruent in the ‘attend auditory’ condition was significant in both left (t_18_ = 3.6, p = 0.002) and right (t_18_ = 3.9, p = 0.001) SMG ROIs. For completeness, there were no significant effects in either ROI in the ‘attend visual’ condition (left: t_18_ = 1.1, p = 0.3; right: t_18_ = 0.8, p = 0.5), matching the absence of activations within the cortical mask.

### Correlation analyses

Finally, we tested whether pseudoword-shape activation magnitudes correlated, across participants, with the magnitude of individual congruency effects derived from in-scanner RTs. The congruency effect computation used the formula (RT_i_ – RT_c_) / (RT_i_ + RT_c_), where RT_i_ and RT_c_ represent RT for incongruent and congruent trials, respectively. We also tested for correlations with scores on the verbal sub-scale of the OSIVQ (Blazhenkova and Kozhevnikov, 2009) and with individual preferences for object compared to spatial imagery by reference to the OSdiff score (see Methods). To avoid circularity, we conducted these correlation tests in the whole brain, i.e. independently of the activations, and in a similarly stringent manner, by setting a strict voxel-wise threshold of p < 0.001 within the cortical mask before applying cluster correction (corrected p < 0.05).

In the ‘attend visual’ condition, activation magnitudes for the C > I contrast were negatively correlated with the magnitude of the in-scanner RT congruency effects at a focus in the left SMG (Table 3a; Figure 8); this focus overlapped with the left SMG focus in the language control task. Additionally, a single voxel in the right SFS, contiguous with right SFS activation on the language control task (Table 3b; Figure 8), had a C > I activation magnitude that was positively correlated with verbal scores from the OSIVQ.

**Figure 8.**
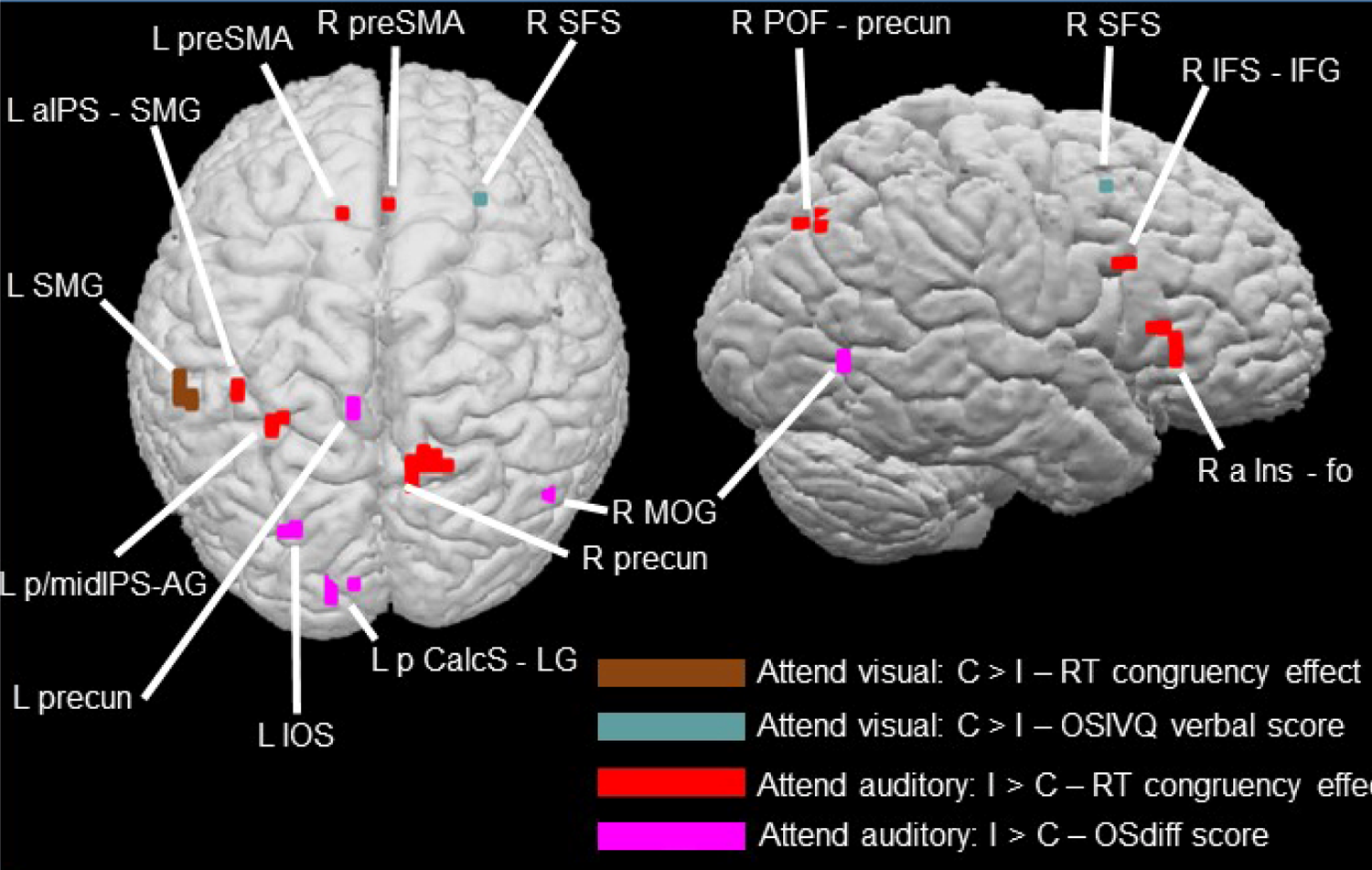
Cortical regions showing correlations between activation magnitudes for the C > I contrast in the ‘attend visual’ condition and the magnitude of the RT congruency effect (brown: Table 3a) and scores on the verbal sub-scale of the OSIVQ (blue-gray: Table 3b); and between activation magnitudes for the I > C contrast in the ‘attend auditory condition and the magnitude of the RT congruency effect (red: Table 3c) and OSdiff scores (magenta: Table 3d) – higher OSdiff scores indicate stronger preference for object, rather than spatial, imagery. See notes to Table 3 for relationships to pseudoword-shape incongruency regions and independent functional contrasts.

**Table 3.**
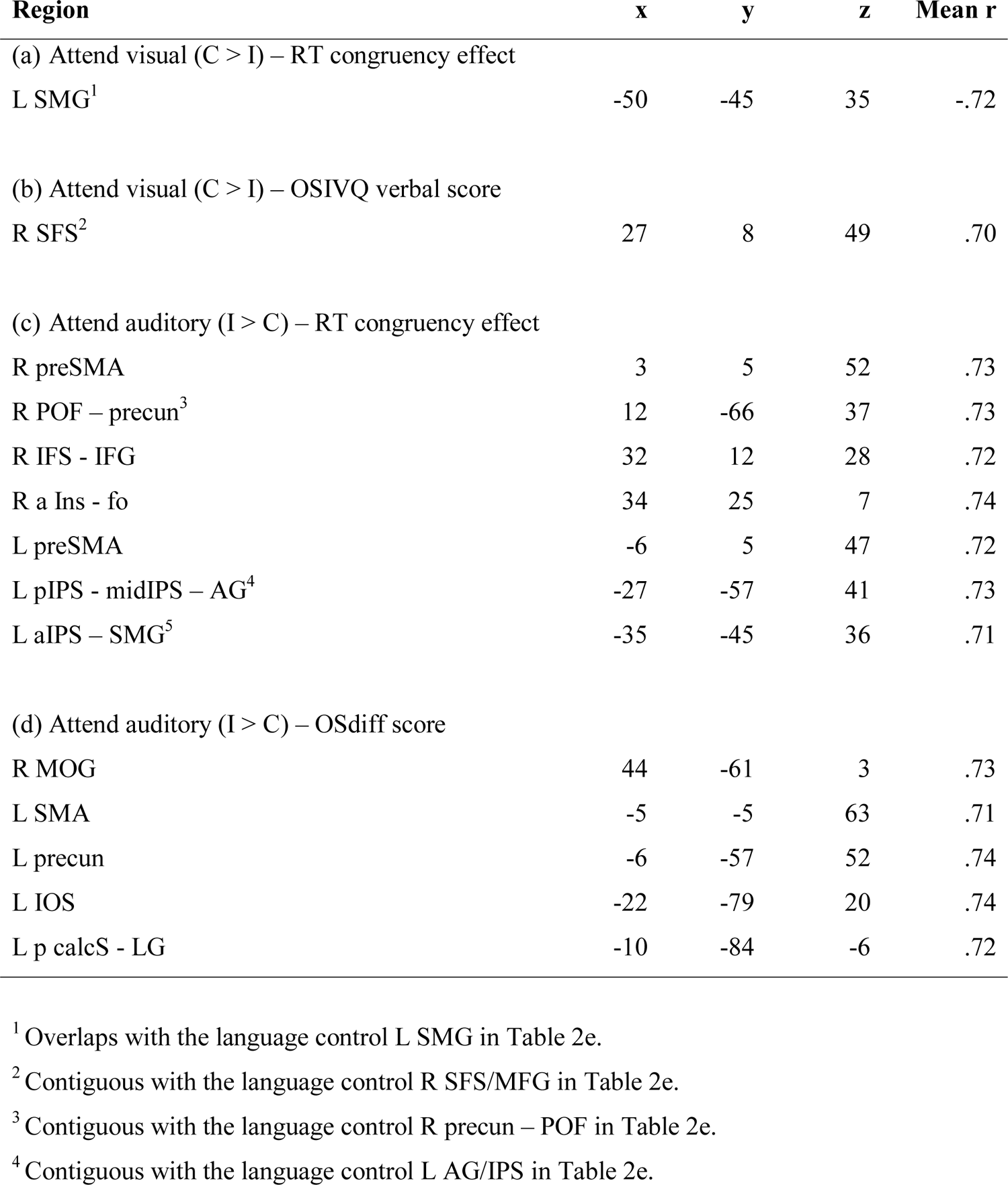
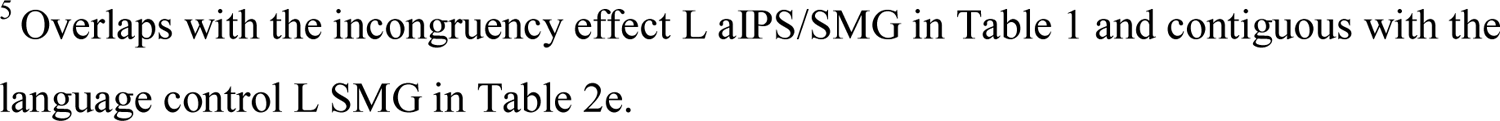
Correlations within cortical mask (r-map threshold p < 0.001, cluster-corrected p < 0.05 (except (b), FDR corrected q < 0.05), cluster thresholds = (a, c) 5 voxels, (d) 4 voxels; x,y,z, Talairach coordinates for centers of gravity.

| **Region** | **x** | **y** | **z** | **Mean r** |
| --- | --- | --- | --- | --- |
| (a) Attend visual (C > I) – RT congruency effect |  |  |  |  |
| L SMG<sup>1</sup> | -50 | -45 | 35 | -.72 |
| (b) Attend visual (C > I) – OSIVQ verbal score |  |  |  |  |
| R SFS<sup>2</sup> | 27 | 8 | 49 | .70 |
| (c) Attend auditory (I > C) – RT congruency effect |  |  |  |  |
| R preSMA | 3 | 5 | 52 | .73 |
| R POF – precun<sup>3</sup> | 12 | -66 | 37 | .73 |
| R IFS - IFG | 32 | 12 | 28 | .72 |
| R a Ins - fo | 34 | 25 | 7 | .74 |
| L preSMA | -6 | 5 | 47 | .72 |
| L pIPS - midIPS – AG<sup>4</sup> | -27 | -57 | 41 | .73 |
| L aIPS – SMG<sup>5</sup> | -35 | -45 | 36 | .71 |
| (d) Attend auditory (I > C) – OSdiff score |  |  |  |  |
| R MOG | 44 | -61 | 3 | .73 |
| L SMA | -5 | -5 | 63 | .71 |
| L precun | -6 | -57 | 52 | .74 |
| L IOS | -22 | -79 | 20 | .74 |
| L p calcS - LG | -10 | -84 | -6 | .72 |

<sup>1</sup> Overlaps with the language control L SMG in Table 2e.
<sup>2</sup> Contiguous with the language control R SFS/MFG in Table 2e.
<sup>3</sup> Contiguous with the language control R precun – POF in Table 2e.
<sup>4</sup> Contiguous with the language control L AG/IPS in Table 2e.
<sup>5</sup> Overlaps with the incongruency effect L aIPS/SMG in Table 1 and contiguous with the language control L SMG in Table 2e.

In the ‘attend auditory’ condition, several foci showed strong positive correlations between activation magnitudes for the I > C contrast and the magnitude of the in-scanner RT congruency effects (Table 3c; Figure 8), overlapping with the ‘attend auditory’ incongruency region in the left aIPS/SMG and close to the left mid-IPS incongruency region. Among the regions identified on the independent localization tasks, correlation foci were contiguous with pseudoword > sentence activations in the right precuneus and the left AG, IPS, and SMG, and close to the language control activation in the right IFS/IFG. A single correlation focus was close to the right SPG activation from the magnitude task. There were also foci at which the magnitude of the BOLD incongruency effect (I > C contrast) was positively correlated with preference for visual object imagery (Table 3d; Figure 8), i.e., magnitudes increased with increasing preference for object relative to spatial imagery. These correlated regions were primarily in the left precuneus, and visual cortical areas: the IOS, the posterior calcarine sulcus extending into the lingual gyrus, and the right MOG.

## DISCUSSION

### Comparison of present study with prior studies

To our knowledge, the present study is the first attempt to investigate, in a principled way, various explanations of the neural basis of *how* sound-symbolic associations are processed, as opposed to merely the neural loci *at* which they are processed. Previous studies have employed sound-symbolic words and pseudowords relating to a range of different domains (Ković et al., 2010; Revill et al., 2014; Sučević et al., 2015; Lockwood et al., 2016; Peiffer-Smadja and Cohen, 2019), but the neural processes involved may differ depending on the particular sound-symbolic association (Sidhu and Pexman, 2018). To avoid this problem, we concentrated on the specific association between auditory pseudowords and visual shapes, and tested the evidence for competing explanations. In addition, we used a post-scan behavioral task to verify that participants did, in fact, make the intended sound-shape mapping.

Sound-symbolic associations are a form of crossmodal correspondence. Spence (2011) suggests that crossmodal correspondences might reflect processes originating in multisensory integration, magnitude estimation, or semantic mediation, depending on the particular correspondence. These suggestions closely approximate three of five potential explanations for sound-symbolic associations put forward by Sidhu and Pexman (2018) although their report was published after data acquisition for the present study had been completed. The remaining two proposals – shared properties across modalities or species-general mechanisms – were not testable in the current study but cannot be excluded. Likewise, the design of our study is also open to the possibility of alternative explanations not considered by Spence (2011) or Sidhu and Pexman (2018).

Here, we investigated sound-to-shape mapping between auditory pseudowords and visual shapes by manipulating the congruency of this sound-symbolic correspondence and employing fMRI which offers greater anatomical resolution than the EEG paradigms of some prior studies (e.g., Ković et al., 2010; Sučević et al., 2015; Lockwood et al., 2016). Briefly, we compared the resulting neocortical activations to the results of functional contrasts designed to reflect the neural processes underlying potential explanations. There were no significant activations during attention to visual shape, but during attention to auditory pseudowords we found greater activity for incongruent, compared to congruent, pseudoword-shape pairs in bilateral IPS and SMG, and in the left MFG. On comparison to functional contrasts, we suggest that these results provide no evidence for semantic mediation, and limited evidence for processes involved in multisensory integration or magnitude estimation; instead, we propose that they likely reflect phonological processing and/or multisensory attention. We emphasize, however, that the present study is exploratory and intended to serve as a spur to further research.

### Sound-symbolic incongruency effects

During attention to visual shapes, no activations survived correction in either the C>I contrast or the reverse. Behaviorally, there was no difference in accuracy between congruent and incongruent trials during attention to shapes, and accuracy was just as high as during attention to pseudowords. RTs during attention to shapes were overall faster than during attention to pseudowords and, although they were slower for incongruent, compared to congruent, trials, this difference was 70% smaller than that during attention to pseudowords. This difference in congruency effects depending on the attended modality may arise from timing differences in auditory and visual processing. The visual shapes appeared in their entirety at the start of each trial and remained onscreen for 500 ms. By contrast, auditory pseudowords, perforce, unfold over time, which may have made them more subject to influence of the unattended modality. In other words, during attention to shapes, a participant could see the entire shape immediately and prepare a response almost at once, even before the auditory pseudoword had been completely presented: note that mean response times for visual shapes (overall 474ms, congruent 469ms, incongruent 480ms) were all shorter than the duration of either pseudoword (‘keekay’ 533ms, ‘lohmoh’ 600ms). This suggests that the unattended pseudowords may have had little influence on either neural processing or behavior when participants attended to the visual shapes.

Although several studies indicate that, even when attention is focused on a single modality of a multisensory object, information in an unattended (and task-irrelevant) modality is processed as well (Busse et al., 2005; Driver and Spence, 2000; Miller, 1991; Molholm et al., 2007; Zimmer et al., 2010), a more recent study suggests that intermodal processing is modulated by attention at a relatively late stage (Shrem and Deouell, 2017). Thus, the asymmetry we observed here between attention to visual and auditory stimulus components might reflect this relatively late influence of crossmodal attention, as it might have been more likely to influence the temporally-unfolding pseudowords, which required more time to process in full than the visual shapes.

Additionally, when there is a conflict between visual and auditory sources of information, there may be a processing bias towards the visual modality (Ben-Artzi and Marks, 1995; Molholm et al., 2004). Such a bias could result in auditory information having less of an influence during attention to vision and/or produce a stronger incongruency response when, supposedly unattended, visual information is incongruent with an attended auditory stream.

During attention to auditory pseudowords, we observed greater cortical activity for incongruent, compared to congruent, pseudoword-shape pairs, which may seem puzzling at first glance. But it is not clear, *a priori*, whether the congruent or incongruent audiovisual condition would evoke the greater activity. On each trial in the imaging experiment of the present study, participants were only required to identify which pseudoword or shape was presented, depending on the attended modality; thus, the congruency or incongruency of the pseudoword/shape pair was only processed implicitly. Similarly, in Peiffer-Smadja and Cohen (2019), participants responded to ‘oddball’ trials and not to pseudowords or shapes at all; thus, here too, sound-symbolic (in)congruency was only processed incidentally and incongruency effects were found. By contrast, a preliminary report of another study from our group, using an *explicit* match/mismatch decision on each trial, found that a focus in the right caudate nucleus exhibited greater activity for congruent than incongruent stimuli, whereas a left precentral focus showed the opposite preference (Barany et al., 2021). Furthermore, a recent study of Japanese mimetic words relating to tactile hardness/softness, in which participants explicitly judged congruency between tactile stimuli and auditory mimetic words on every trial, also found activations for congruency relative to incongruency (Kitada et al., 2021). Thus, effects favoring congruent over incongruent conditions have so far been found only when participants rendered congruency judgments during scanning, implying that this kind of task may be a prerequisite for finding such effects. Effects in the opposite direction, i.e., favoring incongruent over congruent conditions, on the other hand, have been reported as the only or dominant effect independently by multiple studies of sound-to-shape mapping and across different experimental designs and tasks: how are these to be interpreted?

One view of congruency manipulations as a test of multisensory integration is that multiple sensory inputs are available to brain regions involved in integration, and that such regions respond more to congruent than incongruent multisensory inputs (Noppeney, 2012). However, Noppeney (2012) also argued that the brain might attempt, unsuccessfully, to integrate incongruent multisensory inputs before rejecting them as not belonging together. If so, it would not be unexpected to find greater activity for incongruent than congruent stimuli as a result of the more effortful processing involved. In fact, separate previous studies, of audiovisual integration (Hein et al., 2007) and crossmodal visual priming of auditory targets (Noppeney et al., 2008), have both found incongruency effects – importantly, these were not studies of sound symbolism. Incongruent priming effects appeared to be meaningful in that they were selective for spoken words in L STS and for environmental sounds in L AG, while L IFS and prefrontal cortex showed incongruency effects for both (Noppeney et al., 2008).

One way in which incongruency effects might arise is because congruent pairings represent a type of intersensory redundancy, deriving from, or referring to the same semantic source or object and thus requiring fewer processing resources (Bahrick et al., 2004; Shams and Seitz, 2008). If sound symbolism is based in a multisensory system that responds to intersensory redundancy, this could account for the incongruency effects observed in our study and others (Peiffer-Smadja and Cohen, 2019; Barany et al., 2021), possibly due to the added burden of attempting to integrate incongruent or otherwise mismatched sensory inputs (Noppeney et al., 2008). In the present study, it is therefore interesting to note that the R aIPS and R SMG-poCS foci showing greater activity for mismatched than matched pseudoword-shape pairs (Figure 6A) both overlapped with regions more sensitive to audiovisual asynchrony than synchrony (Figure 6C); i.e., both foci were processing items that did not ‘go together’ in some way.

### Relationship of incongruency effects to independent functional contrasts

#### Semantic and phonological processes

The regions identified on the language task showed no overlap at all with the observed activations due to incongruent, compared to congruent, pairings of auditory pseudowords and visual shapes. Thus, of the *a priori* explanations considered in the Introduction, the possibility of semantic mediation, i.e. correspondence between meanings that might be implicit in the pseudowords (e.g., by reference to relevant words with similar phonological content) and the shapes that are associated, appears least likely. However, the incongruency-related region in the left SMG was also more active for pseudowords than complete sentences in the language task. In addition, foci demonstrating correlations of neural incongruency effects with in-scanner RT incongruency effects during the ‘attend auditory’ condition, in the right precuneus and right inferior frontal cortex, were adjacent to or near areas with a preference for pseudowords over sentences.

The premise of sound symbolism is that the sound structure of words mimics or relates to some aspect of what they represent – either explicitly, and within-modally, in the auditory domain as in onomatopoeia (Catricalà and Guidi, 2015; Schmidtke et al., 2014); or by analogy, and crossmodally, as for ideophones or mimetics which can also refer to non-auditory meanings (Akita and Tsujimura, 2016; Kita, 1997). For example, the repetitive sound of the Japanese mimetic term ‘kirakira’ may be considered analogous to the flickering quality of light it describes. It is thus interesting that we found a left-lateralized region in the SMG that was common to both the pseudoword-shape incongruency effect and the pseudoword condition of the language task, even though pseudowords were presented auditorily in the sound-symbolic task and visually in the language control task. Given that reading pseudowords likely depends on phonological processing (Fedorenko et al., 2010) whereas reading sentences also involves syntactic and semantic processing, this suggests that phonological processes may underlie this commonality. Both the left and right SMG are involved in phonological processing (Hartwigsen et al., 2010; Oberhuber et al., 2016), and the left SMG focus has previously been implicated in the phonological processing of visually presented words (Price et al., 1997; Wilson et al., 2011). In order to test the phonology hypothesis further, we conducted an ROI analysis of bilateral SMG foci derived from an earlier study (Hartwigsen et al., 2010), i.e., independently of our findings. This ROI analysis showed incongruency effects in bilateral SMG in the ‘attend auditory’, but not visual, condition. Based on these findings, we suggest that phonological processing could contribute to the pseudoword-shape correspondence. Since the phonological account was not originally hypothesized, and hence the language task was not designed to test this, we cannot currently rule out alternative explanations, so this idea requires further investigation.

There are many aspects to phonology, e.g., syllable structure, stress, and prosody, and while the pseudowords > sentences contrast may reflect phonological processing, it likely involves multiple aspects that need to be disentangled. Although several studies suggest that consonants contribute more than vowels to associations with pointed and rounded shapes (e.g., Nielsen and Rendall, 2011; Ozturk et al., 2013; Fort et al., 2015; but see Styles and Gawne, 2017) and that pointed/rounded shapes are associated with plosive/sonorant consonants (Monaghan et al., 2012; Fort et al., 2015), further work is required to investigate the relative contributions of different phonological and/or phonetic features to particular sound-symbolic associations (e.g. Knoeferle et al., 2017; McCormick et al., 2015). Finally, it should be noted that reading pseudowords (as in the control condition of the language task) involves mapping unfamiliar orthographic patterns^3^ to phonological representations and may thus require more effort than reading complete sentences (Shechter and Share, 2020). Therefore, the overlaps of activations in the language control task with those related to sound symbolism may reflect involvement of the domain-general multiple demand system (Duncan, 2013; Fedorenko et al., 2013), rather than, or in addition to, phonological processes (see below).

The finding that brain regions active during processing of sound-symbolic pseudowords are distinct from the canonical language system (Fedorenko et al., 2010, 2011; Bedny et al., 2011) may offer a potential route for rehabilitation after language impairment, for example in stroke patients. To the extent that sound-symbolic and classical language systems involved in structure and meaning are dissociated, either channel could be exploited to tap into meaningful representations if the other channel showed a deficit. In the same way that hearing a tool sound can remind patients with apraxia how to use that tool (Worthington, 2016), intensive exposure to sound-symbolic words might provide a means of rehabilitating language abilities. In fact, there is preliminary evidence to suggest that patients with aphasia process sound-symbolic words better than more arbitrary word-meaning mappings (Meteyard et al., 2015). This possibility is worth exploring in further research.

#### Multisensory integration processes

As described in the Results, the multisensory task did not show activity in superior temporal cortex, the region most commonly activated in prior studies employing audiovisual (a)synchrony to demonstrate multisensory integration. Most previous studies employed audiovisual speech (van Atteveldt et al., 2007; Stevenson et al., 2010; Noesselt et al., 2012; Erickson et al., 2014) or combinations of familiar environmental sounds and images (e.g., Hein et al., 2007; Noppeney et al., 2008). Since both these stimulus types would be expected to trigger some form of semantic processing, we avoided them given that semantic mediation might have been covered separately by the language task as a potential explanation for the correspondence between spoken pseudowords and visual shapes (although an EEG study suggests similar multisensory integration effects for both speech and non-speech stimuli [Stekelenburg and Vroomen, 2007]).

Additionally, asynchronous audiovisual stimuli are more likely to be perceived as synchronous if the visual element precedes the auditory element (V-A) compared to the reverse (A-V) (e.g., Powers et al., 2009, using simple perceptual stimuli as in the present study; and see Bhat et al., 2015, for the same effect using audiovisual speech stimuli). Since there were equal numbers of V-A and A-V trials in the asynchronous condition, its effectiveness may have been reduced, if some of the V-A trials were actually perceived as synchronous. In order to compensate for this and to test the multisensory integration hypothesis independently of the multisensory synchrony task, we conducted a further ROI analysis of foci in the STS, a classical multisensory region, derived from a previous study involving non-verbal, but semantically related, stimuli (Werner and Noppeney, 2010), but this showed no significant sound-symbolic effects in either of the attended modalities.

Nonetheless, pseudoword-shape incongruency-related activations in the right aIPS and SMG overlapped with areas active during the asynchronous condition of the multisensory task. As noted above, whether mismatched pseudoword-shape pairs (Figure 6A) or auditory and visual stimuli that were presented asynchronously (Figure 6C), both foci appear to be involved in processing items that did not ‘go together’ in some way. The right aIPS incongruency site is close to several IPS foci activated during processing of audiovisual spatial congruency (Sestieri et al., 2006). The IPS portions of our parietal incongruency regions also overlap with regions implicated in multisensory attention during difficult tasks (Regenbogen et al., 2018). This may be relevant to the contention that the brain is trying harder to integrate these inputs when they are incongruent (Noppeney, 2012; see also ‘Incongruency Effects and Attention’ below). The left MFG incongruency region likely overlaps with a region in inferior frontal cortex responding to audiovisual incongruency for familiar environmental stimuli (Hein et al., 2007) and incongruency between visual primes and auditory targets, whether environmental sounds or spoken words (Noppeney et al., 2008). Since the relevant effects were evoked by multiple types of stimulus incongruency, either of content or of timing, the most plausible explanation for these overlaps is the attention-driven effects cited above, perhaps due to unsuccessful attempts to integrate the audiovisual stimuli.

Although foci that responded to incongruent pseudoword/shape pairs also responded to temporal asynchrony, a limitation of the multisensory functional contrast that was intended to broadly reflect multisensory integration is that it may be overly specific to integration arising from temporal co-occurrence, which may not be important in assigning pseudowords to spatial entities like shapes. Certainly, integration can also arise from other types of congruency, such as spatial co-location (Spence, 2013). However, whether spatial co-location is important for integration may be modality- and task-dependent, being less important for some audiovisual (e.g., Regan and Spekreisje, 1977) than visuotactile tasks (e.g., Sambo and Forster, 2009), perhaps especially if, as here, the task is simply to identify the stimulus (Spence, 2013). As noted by Spence (2013, footnote b), auditory stimuli presented via headphones and visual stimuli presented on a screen are different spatial locations and it may be difficult to determine the role of spatial matching in such a set-up. However, participants were both faster and more accurate for congruent than incongruent trials in the ‘attend auditory’ condition, indicating that the spatially different sources of auditory pseudowords and visual shapes had little effect (in the ‘attend visual’ condition, participants were faster, but not significantly differentially accurate, for congruent trials).

Finally, we avoided audiovisual speech or environmental sound and images in the multisensory task on the grounds that these might trigger semantic processes that might also have been reflected in the language task to some extent. However, since sound symbolism implicitly assumes that ‘keekay’/‘lohmoh’ and the pointed/rounded shapes are related, an argument could be made that a multisensory task targeting semantic (in)congruency, either audiovisual speech or environmental stimuli, might have been employed as an independent test of congruency itself. In this respect, it is interesting that we find an incongruency region in the left MFG (and outside the classic language network) that potentially overlaps with that found by Hein et al. (2007) as responsive to semantically incongruent environmental audiovisual pairs. Ultimately, further research is needed to more fully explore the relationship between multisensory integration, semantic processes, and sound symbolism.

#### Magnitude estimation processes

In the present study, the right SMG incongruency region overlapped with activation during magnitude estimation and was contiguous with the right aIPS magnitude-sensitive region. The IPS is a classical locus for magnitude processing (Sathian et al., 1999; Eger et al., 2003; Pinel et al., 2004; Piazza et al., 2004, 2007; Sokolowski et al., 2017) while the SMG is among regions showing adaptation to magnitude (Piazza et al., 2007) or involved in detecting changes in numerosity (Piazza et al., 2004). Additionally, part of the right POF-precuneus region, whose activation magnitude correlated with RTs during attention to auditory stimuli in the scanner, was close to the right SPG activation evoked by magnitude estimation in the independent localization contrast – this region has been implicated in counting visual objects (Sathian et al., 1999). The two pseudowords used in this experiment were drawn, like the shapes, from a much larger set which were empirically rated as sounding more or less rounded or pointed (McCormick et al., 2015). It follows that participants had no problem in applying magnitude estimation (of roundedness or pointedness) to pseudowords. As noted earlier, sensory features often relate to polar dimensions of magnitude where one end is ‘more than’ the other (Smith and Sera, 1992), for example brighter vs. darker or louder vs. quieter, and a contrast targeting such sensory dimensions may have been relevant to the dimensions of pointedness and roundedness. We expected that magnitude processing would be reflected in congruent and incongruent trials because congruent trials involve stimuli occupying similar positions on a continuum whilst incongruent trials involve stimuli occupying different positions. It remains possible that the incongruency effects do indeed reflect magnitude processing because there are different magnitudes involved. However, the role of magnitude processing might become clearer in a parametric setting, using a range of pseudowords and shapes and thus sampling at multiple intervals along the rounded/pointed continuum (whilst accepting that the continuum might not be consistently mapped across individuals). Despite extensive overlap between brain regions involved in processing number- and sensory-based magnitude (Pinel et al., 2004), caution is indicated by recent challenges to the idea of a domain-general magnitude system (Anobile et al., 2018; Pitt and Casasanto, 2018; but see Holmes et al., 2019) which has been posited as an explanation for crossmodal correspondences (Lourenco and Longo, 2011; Spence, 2011; Sidhu and Pexman, 2018). Additionally, while we cannot rule out magnitude estimation as a possible explanation for the observed sound-symbolic incongruency effects, the regions of overlap between the magnitude estimation task and the incongruency effects are also implicated in multisensory attentional processes, which may ultimately turn out to be more important (see below).

### Incongruency effects and attention

No brain regions showed congruency effects in either attended modality in the present study, which was also the case in the study of Peiffer-Smadja and Cohen (2019). However, when participants were attending to the auditory pseudowords, several neocortical regions showed greater activity for incongruent, compared to congruent, pairings with visual shapes. These areas were more widespread than the dorsolateral prefrontal regions found by Peiffer-Smadja and Cohen (2019), who used a task requiring only occasional responses (see below). We did not find incongruency effects when participants attended to the visual shapes, consistent with a behavioral congruency effect being absent in the ‘attend visual’ runs when accuracy was the dependent measure, and being significantly smaller in the ‘attend visual’ than the ‘attend auditory’ runs when RT was the dependent measure. These differences as a function of the attended modality may stem from greater unfamiliarity of auditory pseudowords relative to two-dimensional visual shapes. Alternatively, they might reflect timing differences in processing pseudowords compared to visual shapes, since the pseudowords unfold over time, whereas the visual shapes are in evidence from the start of each trial. Consequently, there may be more of an incongruency effect in the ‘attend auditory’ condition because the concurrent visual stimulus could trigger particular representations immediately, whereas in the ‘attend visual’ condition, the concurrent pseudoword may still be playing as the participant prepares a response to the visual stimulus.

The regions demonstrating activations attributable to incongruency were in various parts of parietal cortex bilaterally, and in the left MFG. The left MFG activation likely overlapped with an extensive zone of frontal cortical sensitivity to pseudoword-shape incongruency in the study of Peiffer-Smadja and Cohen (2019), although this was associated with greater deactivation for congruent than incongruent stimuli in that study and, as noted earlier, deactivations are hard to interpret (see Footnote 2). Furthermore, since the task in that study was to detect rare occurrences of visual crosses and auditory beeps, the sound-symbolic pseudoword and shape stimuli were actually irrelevant to the main detection task; thus the frontal deactivation to mismatched pseudoword-shape pairs may well have arisen because these were a distraction from the explicit detection task (Peiffer-Smadja and Cohen, 2019).

An incongruency effect of the type reported here might be enhanced when, as in the present study, participants selectively attend to one modality while ignoring the other (Noppeney, 2012). For example, when participants attended auditory targets after passively viewing visual primes, greater activity was elicited in several cortical areas (including one that likely overlaps with our left MFG incongruency region) when primes and targets were incongruent compared to congruent (Noppeney et al., 2008). Our left MFG incongruency region likely overlapped with a focus displaying greater activity for audiovisual stimuli that were familiar but semantically incongruent, compared to unfamiliar arbitrary pairings of abstract images and sounds (Hein et al., 2007). This region also corresponds to part of the domain-general ‘multiple demand’ attentional system described by Duncan (2013). In the ‘attend auditory’ incongruent condition, then, what may be happening is that the brain attempts to integrate a stimulus pair but instead detects a mismatch between a rounded pseudoword and a pointed shape or vice versa. On this reasoning, the basis of the observed incongruency-related activations might be greater attentional demand, rather than multisensory integration, during discrimination of (unfamiliar) auditory pseudowords in the presence of incongruent, compared to congruent, visual shapes. Consistent with this, the incongruency region in the right SMG is close to a focus exhibiting a preference for novel over familiar stimuli across auditory, visual, and tactile modalities (Downar et al., 2002), and to one in which activation was stronger for unfamiliar abstract audiovisual stimuli compared to familiar, but semantically incongruent, audiovisual stimuli (Hein et al., 2007). Furthermore, the incongruency regions found here in bilateral aIPS and left SPG correspond to regions active during top-down attentional processing when visual targets were cued by semantically incongruent audiovisual stimuli (Mastroberardino et al., 2015), and to bilateral IPS foci (both of which overlap with our incongruency regions) shown to mediate audiovisual interaction during difficult processing of degraded stimuli (Regenbogen et al., 2018). This attentional explanation is especially likely for the subset of incongruency regions, comprising foci in the left IPS/SMG that also exhibited correlations between the magnitudes of these neural incongruency effects and the corresponding RT congruency effects during scanning. We acknowledge that these considerations are subject to the limitations of reverse inference, although the limitations tend to be mitigated by our use of task-related reverse inference (Hutzler et al., 2014; McCormick et al., 2018). We suggest that the role of multisensory attention in relation to sound-symbolic crossmodal correspondences merits further research.

### Relationship to visual imagery

Across participants, increasing preference for object as opposed to spatial imagery was strongly positively correlated with activation magnitudes for the incongruent > congruent contrast in the ‘attend auditory’ condition. Many of these correlated regions were in visual cortex (in the left IOS, posterior calcarine sulcus/lingual gyrus, and the right MOG), and one was in the left precuneus. This may reflect a greater tendency for object imagers to visualize shapes when associating these with pseudowords. The precuneus (Cavanna and Trimble, 2006), calcarine sulcus, lingual gyrus and MOG (Ganis et al., 2004) are all involved in visual imagery. Another possibility is that, since object imagers tend to integrate structural properties like shape with surface properties like texture (Lacey et al., 2011), the phonetic properties of the pseudowords, for example whether or not consonants are voiced or vowels are rounded, are more strongly bound to the visualized pointed and rounded shapes in these individuals. Interestingly, in contrast to the relatively widespread correlations with imagery preferences, there was little evidence that preference for verbal processing was connected to sound-symbolic associations since correlation of activation magnitudes with OSIVQ verbal scores was limited to a single voxel in the right SFS. Further work is required to investigate the potential for individual differences since these might underlie the ability to either guess the meaning of sound-symbolic words in unknown foreign languages (Kunihira, 1971), or to learn such associations (Nygaard et al., 2009; Revill et al., 2018).

### Limitations

We note some limitations of the current study. Firstly, as noted above, alternative functional contrasts might have been adopted for both the magnitude and multisensory integration tasks. The magnitude task essentially involved a numerosity judgment whereas the pseudowords and shapes were derived from a rounded-pointed continuum. The domain-general magnitude system suggested as a potential basis for crossmodal correspondences (Spence, 2011) and some instances of sound symbolism (Sidhu and Pexman, 2018) would be expected to be involved in both numerosity and attribute magnitude or intensity (see Walsh, 2003). But it is possible that a more sensory-based magnitude continuum, e.g., lighter-darker (see Pinel et al., 2004), would have been more akin to the sensory pointed/rounded continuum of the pseudowords and shape stimuli. For the multisensory integration task, we avoided speech and environmental stimuli because semantic processing was covered by the language task. But the language task did not address semantic (in)congruency directly, which is arguably part of what the pseudoword-shape pairs reflected. Thus, a functional contrast addressing semantic (in)congruency might have captured a more specific aspect of the sound-symbolic pairs. We should also note that, in the sound-symbolic task, all conditions were multisensory; thus, we were unable to compare activity in these conditions to that in unisensory auditory and visual conditions.

The pseudowords were spoken by an adult female and we did not vary the speaker (e.g. to include male or children’s voices). While including multiple speakers might be desirable in studies of speech perception, it is common for studies of sound symbolism to restrict stimuli to a single speaker voice or to use a synthetic voice. One reason this is important is that different voices could be associated with particular stimuli independently of the words they are uttering. For example, a child’s voice is higher pitched than an adult’s, and female voices are typically higher pitched than male voices; higher pitch in either case could be associated with more pointed shapes. Using a single speaker as we did allows us to avoid these confounds and examine the role of phonological structure of the pseudowords of interest. Nonetheless, we are typically exposed to a variety of voices in normal life and it would thus be desirable to incorporate multiple speakers, balancing the number of utterances across male and female child and adult speakers. Additionally, one could manipulate the factor of speaker variability and investigate whether responses to pointed shape are greater for higher-pitched children’s voices than lower-pitched adult voices and whether responses to rounded shapes are greater for female compared to male voices.

Finally, we only used a single rounded and pointed exemplar pseudoword and shape, which might restrict generalizability of the present findings. However, these stimuli were chosen because they represent extreme values along the rounded-pointed continuum and thus could offer the strongest possible response for congruent/incongruent pseudoword-shape pairs. Critically, sound-symbolic mappings appear to be both graded and relative (Thompson and Estes, 2011); thus, if we had used multiple exemplars, some would likely have been perceived as less rounded or pointed than others, which could raise issues with interpreting congruency. Future work could address this via multivariate analyses in a parametric setting, using a range of pseudowords and shapes and thus sampling at multiple intervals along the rounded/pointed continuum (whilst accepting that the continuum might not be consistently mapped across individuals).

## Conclusions

This study is the first to systematically examine competing explanations underlying the sound-symbolic crossmodal correspondence between auditory pseudowords and visual shapes. We used independent contrasts and ROI analyses to test a number of *a priori* hypotheses for the neural basis of these correspondences. Overall, evidence for semantic mediation was lacking while that for multisensory integration and magnitude estimation was weak, at best. Albeit via task-based reverse inference, multisensory attention emerged as an important potential candidate account, as we also proposed previously for the crossmodal pitch-elevation correspondence (McCormick et al., 2018); we also found evidence consistent with phonological processing as an underlying explanation though this requires more rigorous testing. While this study addressed a tractable set of candidate explanations drawn from Spence (2011) and relevant to Sidhu and Pexman (2018), other possibilities remain to be explored, for example the neural basis of the sensorimotor hypothesis proposed by Ramachandran and Hubbard (2001). Nonetheless, our findings provide a basis for further research, which should seek converging evidence using other methods (for example, multivariate fMRI analyses, repetition suppression fMRI studies, or transcranial magnetic stimulation) and should also address the relative weights of these different processes.

Further, we observed several regions in visual cortex in which activation magnitudes scaled with individual preference for object imagery, thus suggesting a basis for individual differences in processing sound-symbolic associations to be followed up in future research. An important goal for research into crossmodal correspondences in general, as well as sound symbolism in particular, will be to distinguish the roles of high-level cognitive processes, such as phonology and attention, from those of lower-level sensory and perceptual processes.

## Abbreviations: Directional

a: anterior

front: frontal

i: inferior

lat: lateral

m: mid

med: medial

p: posterior

s: superior

v: ventral

## Abbreviations: Anatomical

AG: angular gyrus

AOS: anterior occipital sulcus

calcS: calcarine sulcus

CeS: central sulcus

cingG: cingulate gyrus

cingS: cingulate sulcus

collatS: collateral sulcus

FG: fusiform gyrus

fo: frontal operculum

InfOS: inferior occipital sulcus

Ins: insula

IOG: inferior occipital gyrus

IOS: intra-occipital sulcus

IPS: intraparietal sulcus

ITG: inferior temporal gyrus

ITS: inferior temporal sulcus

LG: lingual gyrus

MFG: middle frontal gyrus

MOG: middle occipital gyrus

OrbG: orbital gyrus

poCG: post central gyrus

po: pars opercularis

poCS: post central sulcus

POF: parieto-occipital fissure

preCS: precentral sulcus

preCG: precentral gyrus

precun: precuneus

preSMA: presupplementary motor area

pt: pars triangularis

SFG: superior frontal gyrus

SFS: superior frontal sulcus

SMG: supramarginal gyrus

SOG: superior occipital gyrus

SPG: superior parietal gyrus

STS: superior temporal sulcus. All other abbreviations are as in the main text.

## ACKNOWLEDGMENTS

We thank Lawrence Barsalou, Justin Bonny, Daniel Dilks, Evelina Fedorenko, Tami Feng, Harold Gouzoules, Sara List, Stella Lourenco and Kate Pirog Revill for their advice and assistance. An earlier version of this manuscript has been released as a pre-print at bioRxiv.org, doi:10.1101/478347 (McCormick et al., 2018a).

## AUTHOR CONTRIBUTIONS

KM, KS, LCN and SL designed the experiment; KM created stimuli; KM and RS performed the research and analyzed data; KM, SL, KS and LCN wrote the paper.

